# Insulin fails to mitigate amyloid pathology but modulates intestinal morphology in the 5xFAD model revealed by synchrotron-based XPCT

**DOI:** 10.64898/2026.10.10.758244

**Authors:** Esra Ayan, Francesca Palermo, Giorgia Cavioli, Sevginur Bostan, Busra Zuleyha Dogan, Kader Kilic, Arife Ahsen Kaplan, Inna Bukreeva, Lorenzo Massimi, Elena Longo, Gulsena Baydas, Ahmet Kati, Emrah Eroğlu, Gürkan Öztürk, Muhammed Ikbal Alp, Alessia Cedola

## Abstract

Intranasal insulin has been investigated as a metabolic intervention in Alzheimer’s disease (AD), but its effects across the brain–gut axis remain unclear. We combined multiscale synchrotron X-ray phase-contrast tomography (XPCT), amyloid-β immunofluorescence, and fecal metaproteomics to examine structural and molecular responses in 5xFAD mice. Twenty-four mice representing wild-type and 5xFAD genotypes, saline and insulin conditions, and terminal ages of 2 and 6 months received insulin at 1 U/mouse/day for 5 weeks. Micro-XPCT visualized plaque-like deposits, nanoXPCT revealed their heterogeneous internal organization, and MOAB-2 labeling provided qualitative support for amyloid pathology. Detectable plaque burden was not lower after treatment: deposits occurred in two of three treated versus none of three untreated 5xFAD mice at 2 months, and treated animals had higher descriptive regional means at 6 months. In the ileum, insulin was associated with lower villus length and crypt depth in 6-month-old 5xFAD mice, whereas no significant differences were detected at 2 months. Fecal profiles retained broad taxonomic overlap but differed in functional annotations, with hydrolase-associated fractions of 81%, 92%, 96%, and 93% across untreated wild-type, treated wild-type, untreated 5xFAD, and treated 5xFAD conditions, respectively, within the selected peptide subset. Thus, lower ileal dimensions coexisted with non-reduced detectable brain plaque burden and a fecal protein profile distinct from untreated wild type. Together, these findings reveal contrasting central and peripheral patterns following intranasal insulin treatment in 5xFAD mice: no detectable reduction in cerebral plaque burden, alongside attenuation of selected ileal structural alterations and distinct fecal metaproteomic profiles. Although exploratory, the findings raise the possibility that intranasal insulin alters amyloid compartmentalization or plaque detectability rather than promoting amyloid clearance, while producing distinct peripheral remodeling.

## 1. Introduction

Alzheimer’s disease (AD) is characterized by progressive cognitive decline associated with amyloid-β (Aβ) deposition, tau pathology, neuroinflammation, synaptic dysfunction, and neuronal loss (Dibattista et al., 2020). Alongside these established features, impaired insulin responsiveness has emerged as a metabolic dimension of AD. Human brain studies have identified abnormalities in insulin and insulin-like growth factor signaling, although their relationship to systemic insulin resistance and disease progression remains complex (Steen et al., 2005; Talbot et al., 2012). These observations have stimulated interest in interventions that enhance central insulin availability without assuming that improved signaling necessarily entails amyloid clearance. Intranasal insulin offers a route for accessing central nervous system pathways while limiting systemic exposure (Craft, 2012; Freiherr et al., 2013). Early clinical studies reported cognitive or biomarker signals in mild cognitive impairment (MCI) and AD, but the larger SNIFF trial did not demonstrate benefit on its primary cognitive endpoint in the principal device cohort (Craft, 2012; Craft et al., 2017, 2020). Recent PET and short-term intervention studies provide further evidence concerning delivery and metabolic modulation rather than definitive therapeutic efficacy (Erichsen et al., 2025; Sai et al., 2025). Preclinical responses are also context-dependent: selected amyloid-related outcomes improved in 3xTg-AD and APPswe/PS1dE9 mice, under protocols differing in age, treatment duration, and pathological readout (Chen et al., 2014; Mao et al., 2016). The effects of insulin on amyloid-associated patology therefore require evaluation within each experimental setting.

AD is increasingly investigated beyond the brain, including through the microbiota–gut– brain axis (Ayan et al., 2023; Palermo et al., 2025). Human studies describe AD-associated microbial differences, while germ-free, antibiotic, and transfer experiments demonstrate that microbial perturbations can influence amyloid or cognitive phenotypes in particular animal models (Grabrucker et al., 2023; Harach et al., 2017; Minter et al., 2017). Microglial and microbial-metabolite pathways provide possible links between intestinal environments and brain pathology (Colombo et al., 2021; Erny et al., 2015). In 5xFAD mice, altered microbial composition and tryptic activity have been reported, and longitudinal and metabolomic studies emphasize the importance of age and experimental context (Brandscheid et al., 2017; Dunham et al., 2022). These findings motivate cross-organ analysis without establishing that every intestinal change drives a corresponding brain response. Structural and molecular approaches address different levels of this relationship.

X-ray phase contrast tomography (XPCT) provides label-free, three-dimensional visualization of weakly absorbing tissues, enabling analysis of brain deposits and intestinal architecture within their volumetric context (Palermo et al., 2025). Intact-block micro-XPCT (µXPCT) can guide subsequent extraction of selected cores for nanoXPCT, while immunofluorescence supplies complementary molecular information. Metaproteomics, in turn, identifies detectable community proteins and can reveal functional differences not apparent from broad taxonomy alone. Protein annotation is nevertheless distinct from measured enzyme activity or metabolic flux (Ayan et al., 2023; Palermo et al., 2025). Here, we investigated how intranasal insulin is associated with changes across three complementary levels of the brain–gut axis in 5xFAD mice: cerebral amyloid pathology, ileal microarchitecture, and fecal microbial protein profiles. We examined brain and ileal specimens at two terminal ages using multiscale XPCT, with MOAB-2 immunofluorescence providing complementary evidence for Aβ pathology and fecal metaproteomics providing taxonomic and functional protein annotations. By integrating these readouts, we asked whether insulin-associated changes occur in parallel across central and peripheral compartments or instead show compartment-specific patterns.

## 2. Results

The experimental design and analytical workflow are summarized in **Figure 1**, **Fig. S1** and **Fig. S2.** Twenty-four mice were distributed among wild-type (Alm), insulin-treated wild-type (Alm-ins), untreated 5xFAD (Alz), and insulin-treated 5xFAD (Alz-ins) conditions at 2 and 6 months, with three animals per genotype–treatment–age group (**Table 1**). The ages represent separate terminal cohorts. Fecal outputs were available as four-condition summaries; their age assignment and individual-animal linkage were not resolved, so they were not used for animal-level cross-organ correlations.

**Figure 1.**
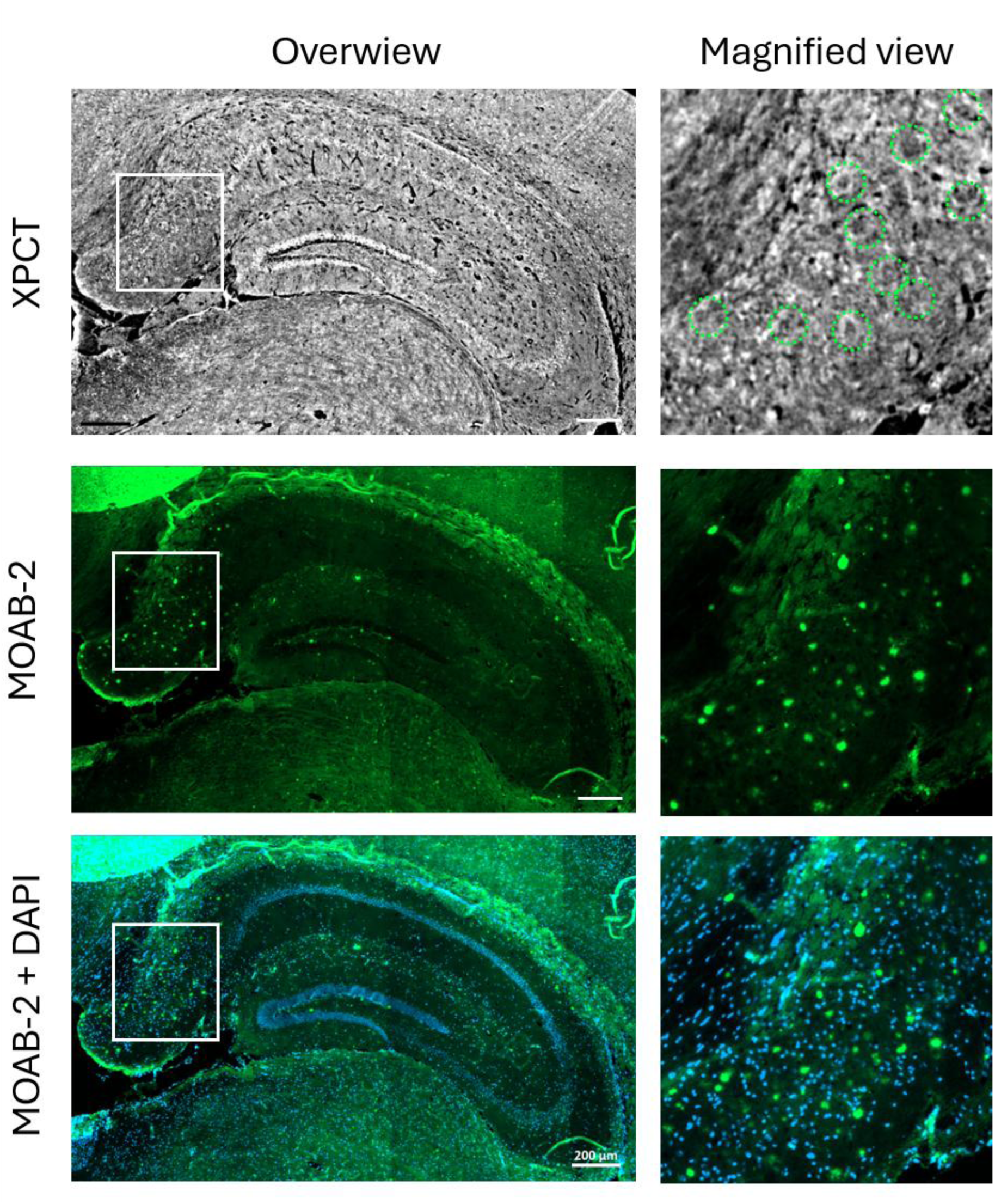
Experimental design and integrated analytical workflow. **(A)** Experimental groups included wild-type C57BL/6 mice (Alm) and 5xFAD mice (Alz), each receiving either intranasal saline or insulin treatment, and analyzed at 2 and 6 months of age. Brain, ileum, and fecal samples were collected for downstream analyses. **(B)** Brain and ileum samples were fixed, paraffin-embedded, and analyzed by micro-X-ray phase-contrast tomography (μXPCT). Brain μXPCT datasets were used for anatomical ROI selection and plaque quantification, and to guide the extraction of 500-µm-diameter cylindrical biopsies for nanoXPCT. Paraffin sections were additionally processed for MOAB-2 immunofluorescence validation. Ileum μXPCT datasets were used for morphometric analysis. Fecal samples were subjected to protein extraction, digestion, and LC–MS/MS-based metaproteomic analysis. **(C)** Main study readouts included amyloid pathology in the brain, intestinal morphology in the ileum, and peptide-based taxonomic and functional profiling of the fecal microbial proteome.

**Table 1.**
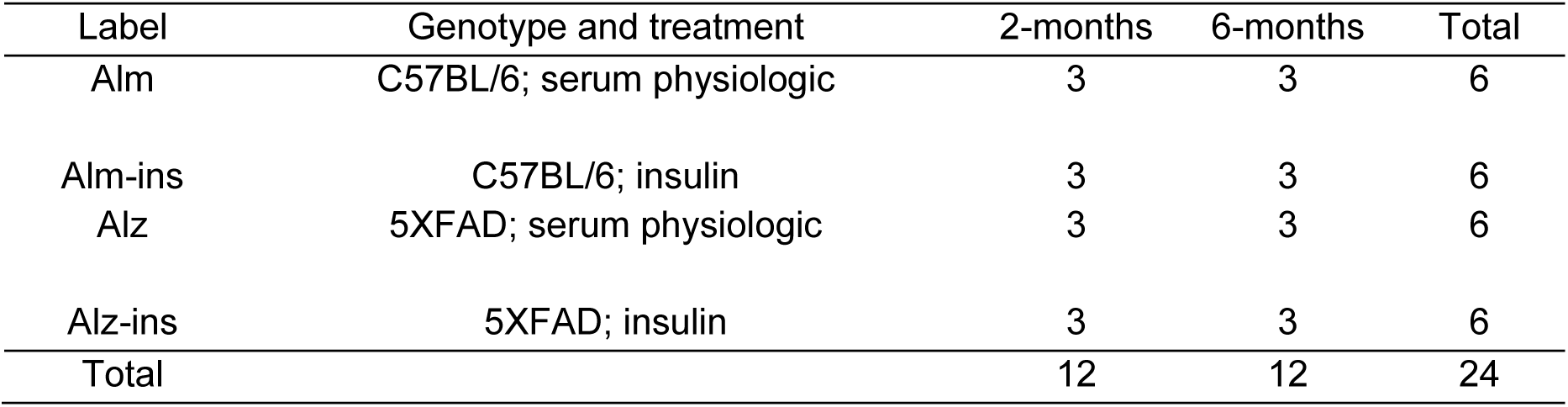
Animal allocation and intranasal treatment. Each age is a separate terminal cohort.

| Label | Genotype and treatment | 2-months | 6-months | Total |
| --- | --- | --- | --- | --- |
| Alm | C57BL/6; serum physiologic | 3 | 3 | 6 |
| Alm-ins | C57BL/6; insulin | 3 | 3 | 6 |
| Alz | 5xFAD; serum physiologic | 3 | 3 | 6 |
| Alz-ins | 5xFAD; insulin | 3 | 3 | 6 |
| Total |  | 12 | 12 | 24 |

### 2.1 Multiscale XPCT characterization of amyloid pathology in 5xFAD mouse brain

Synchrotron-based µXPCT enabled label-free three-dimensional visualization of brain cytoarchitecture and Aβ plaque-like deposits in intact paraffin-embedded 5xFAD mouse brains (**Fig. 2A**). Plaque structures were identified in μXPCT images based on their characteristic morphology and contrast. At higher magnification, they appeared as rounded or spherical features relative to the surrounding brain parenchyma, often displaying heterogeneous internal contrast with a denser central region and a more diffuse peripheral component (**Fig. 2B**). Their morphology and contrast distinguished them from vascular profiles or other anatomical or imaging structures. Plaques were approximately 10–30 µm in dimension, although a quantitative size distribution was not determined. like vessels and capillaries, whose cross section appears similar but their elongated profiles extending across consecutive tomographic slices. The three-dimensional rendering in **Fig. 2D** illustrates the spatial distribution of segmented plaques, shown in yellow, in a selected hippocampal region.

**Figure 2.**
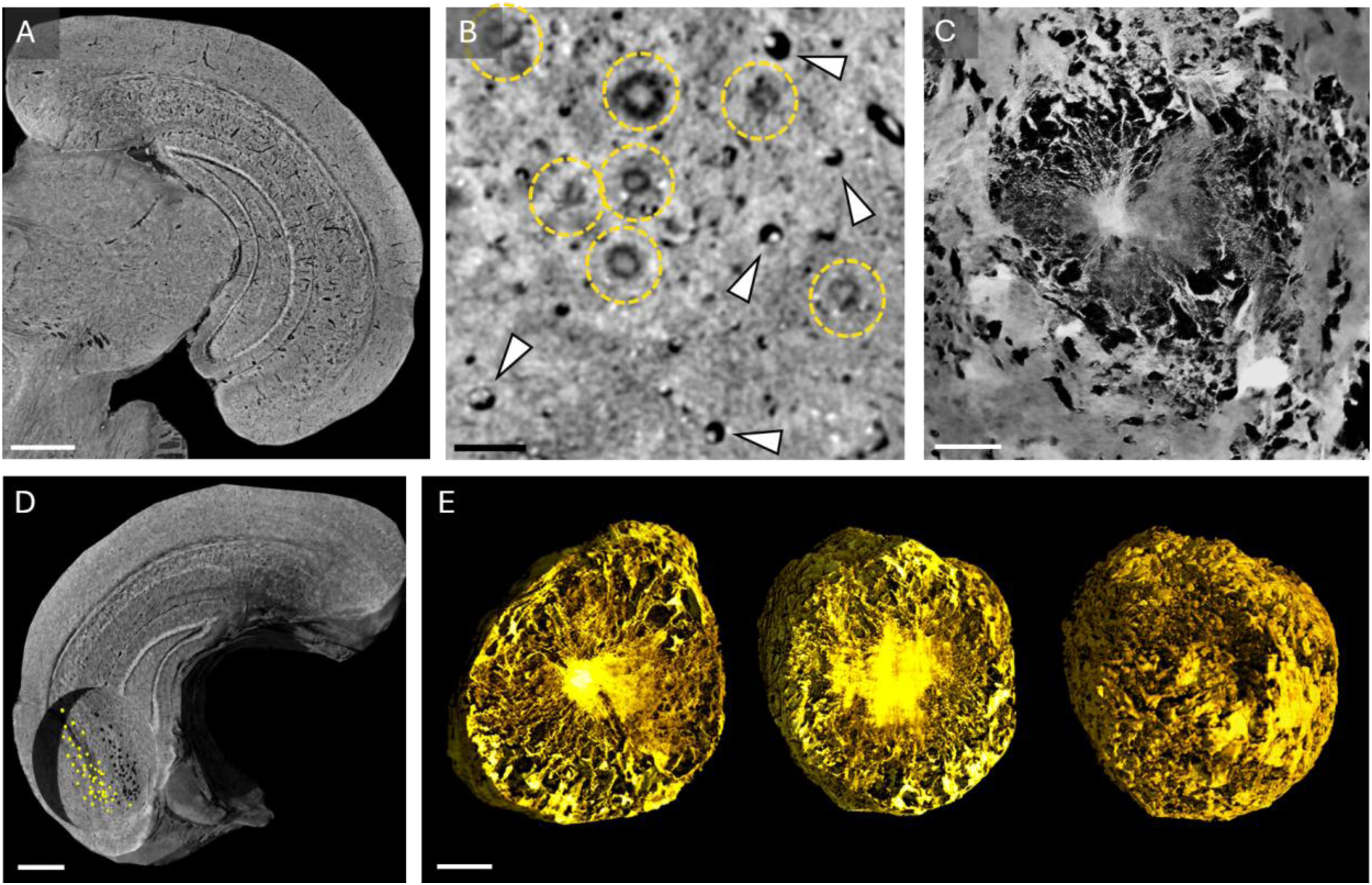
Multiscale μXPCT and nanoXPCT characterization of amyloid-β plaque structures in 5xFAD mouse brains. **(A)** Representative μXPCT slice of a 5xFAD mouse brain, showing a hemispheric section at low magnification. Scale bar: 250 μm. **(B)** High-magnification μXPCT image showing the characteristic morphology of plaque structures, highlighted by yellow dashed contours. The deposits appear as rounded features with heterogeneous internal contrast. White arrowheads indicate vascular profiles that can be distinguished from plaque-like structures based on morphology and contrast. Scale bar: 50 μm. **(C)** Representative nanoXPCT image of an amyloid-β plaque acquired at higher spatial resolution, revealing its heterogeneous internal organization and finer structural features not resolved at the μXPCT scale. Scale bar: 10 μm. **(D)** Three-dimensional rendering of a selected hippocampal region from a 5xFAD mouse. Segmented plaque structures are shown in yellow, illustrating their spatial distribution within the reconstructed volume. Scale bar: 250 μm. **(E)** Three-dimensional renderings of the plaque shown in (C), displayed as two cutaway views to visualize its internal morphology and as the complete reconstructed plaque volume. Scale bar: 10 μm.

NanoXPCT of selected regions permitted a more detailed characterization of plaque morphology, revealing a heterogeneous and non-uniform internal organization of individual plaques that was only partially resolved at the μXPCT scale (**Fig. 2C**). Local contrast variations and a more clearly defined interface between the plaques and the surrounding parenchyma became more evident at the nanoXPCT scale. The Aβ plaques were characterized by a denser central region, suggesting highly compact amyloid material, surrounded by a less compact and composite external layer. The three-dimensional renderings of the plaque shown in (**Fig. 2E**) display two cutaway views to visualize its internal morphology and the complete reconstructed plaque volume

### 2.2. Immunofluorescence supports the Aβ nature of XPCT-detected deposits

To support the amyloid-β nature of the plaque structures identified by XPCT, immunofluorescence staining was performed using the MOAB-2 antibody. In the hippocampal region of a 2-month-old insulin-treated 5xFAD mouse, MOAB-2 staining revealed numerous Aβ-positive deposits in the same anatomical region in which μXPCT-detectable plaque-like structures were observed (**Fig. 3**). MOAB-2-positive deposits were also detected in the hippocampus of a 6-month-old insulin-treated 5xFAD mouse (**Fig. S3**), supporting the presence of established amyloid pathology at the later time point. The correspondence between MOAB-2 immunoreactivity and μXPCT-detected structures was not interpreted on a one-to-one basis. In addition to discrete deposits, the Alexa Fluor 488 channel showed diffuse punctate and vessel-like fluorescence. Signals also visible in the red reference channel or in the no-primary control were considered as autofluorescence or primary-antibody-independent background. In contrast, additional discrete Alexa Fluor 488-positive puncta without a corresponding red-channel signal were considered consistent with Aβ-associated immunoreactivity (**Fig. S4**).

**Figure 3.**
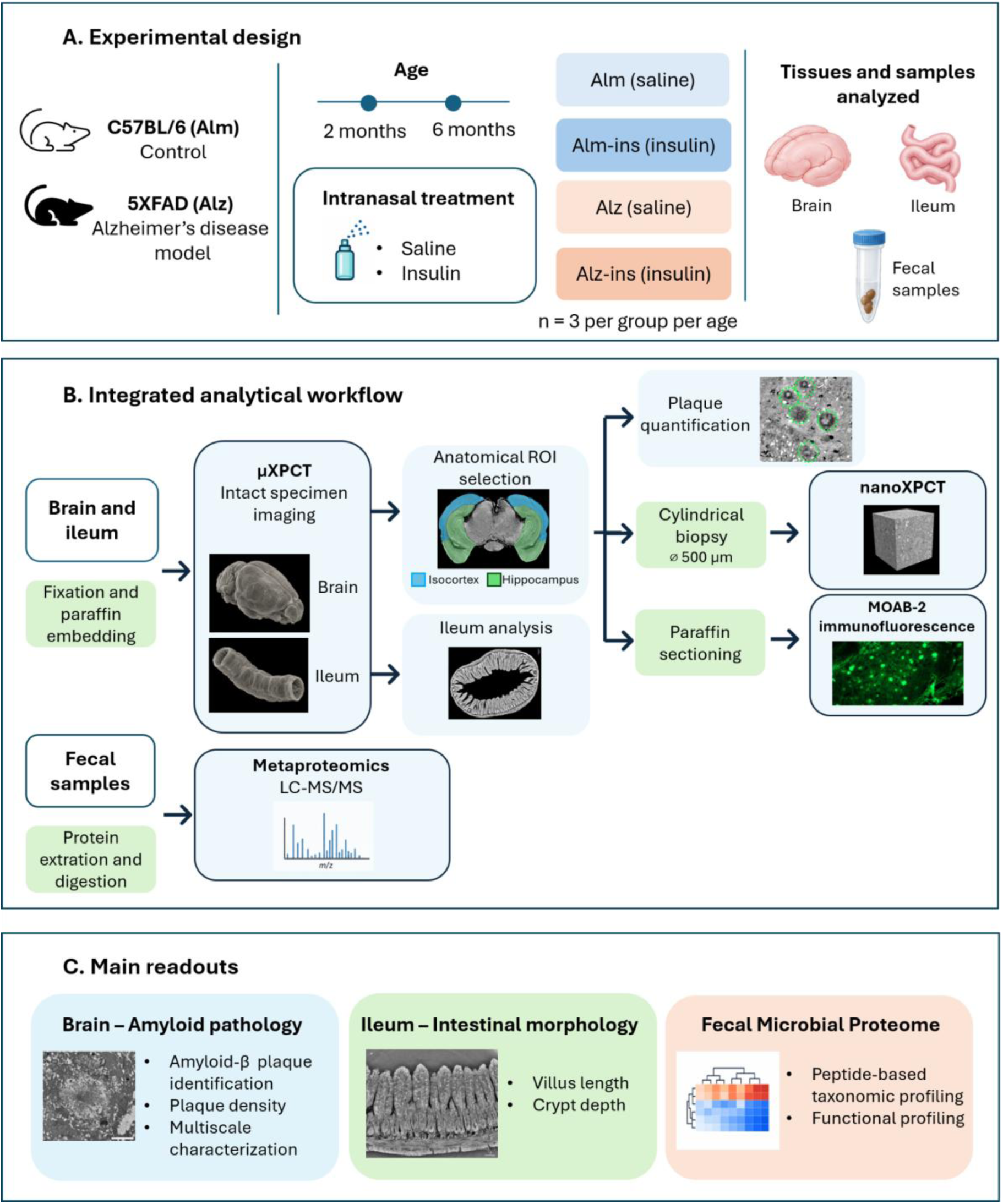
Immunofluorescence support for amyloid-β pathology in a 2-month-old insulin-treated 5xFAD mouse. Representative μXPCT and immunofluorescence images from the hippocampal region of a 2-month-old insulin-treated 5xFAD mouse. The left column shows overview images, while the right column shows magnified views of the boxed regions. The top row shows μXPCT images, with plaque-like structures highlighted by green dashed contours in the magnified view. The middle row shows MOAB-2 immunofluorescence labeling of Aβ-positive deposits, and the bottom row shows MOAB-2 fluorescence merged with DAPI nuclear counterstaining. MOAB-2-positive deposits were observed in the same anatomical region in which μXPCT-detectable plaque-like structures were identified, supporting their association with amyloid-β pathology. The immunofluorescence pattern included a broader range of Aβ-associated signals than the discrete plaque-like structures detectable by μXPCT; therefore, correspondence between the two modalities was not interpreted on a one-to-one basis. Scale bar: 200 μm in overview images.

### 2.3. Insulin is associated with differences in μXPCT-detected plaque burden

Representative μXPCT images showed differences in plaque detectability across age and treatment conditions (**Fig. 4A**). At 2 months, no structures meeting the plaque-detection criteria were identified within the analyzed regions of untreated 5xFAD mice. In contrast, plaque deposits were detected in two of three insulin-treated 5xFAD mice. At 6 months, plagues were detected in two of three untreated and all three insulin-treated 5xFAD mice. Anatomical segmentation delineated the isocortex and hippocampus as regions of interest for subsequent plaque analysis.

**Figure 4.**
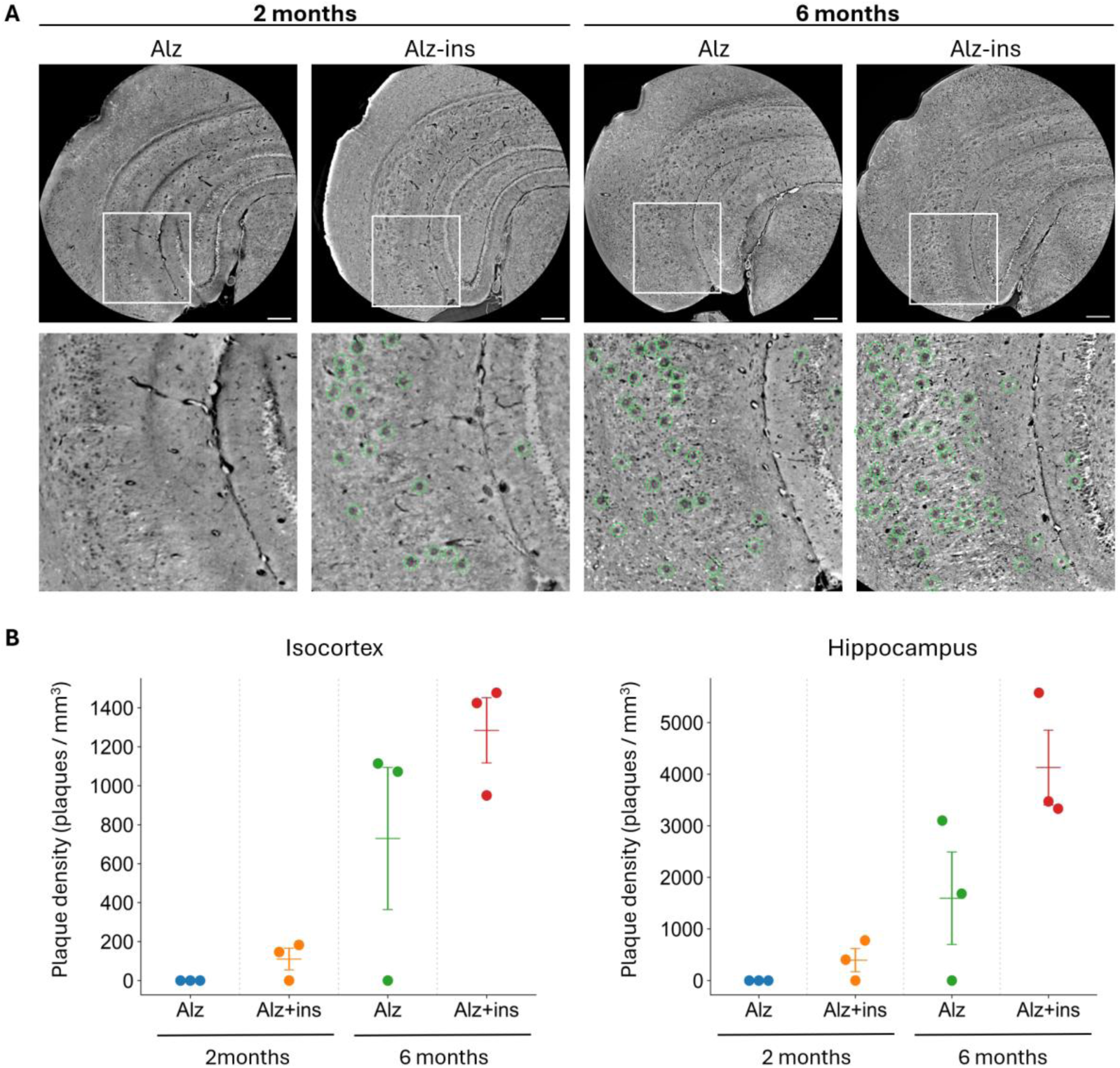
Age- and treatment-associated differences in μXPCT-detected plaque burden in 5xFAD mouse brains. **(A)** Representative μXPCT images from untreated 5xFAD mice (Alz) and insulin-treated 5xFAD mice (Alz + ins) at 2 and 6 months of age. Magnified views of the selected hippocampal regions are shown below the corresponding overview images. Plaque-like structures identified by μXPCT are highlighted by green dashed contours. At 2 months, no readily detectable plaque-like structures were observed in the representative untreated 5xFAD sample, whereas multiple deposits were visible in the insulin-treated animal. At 6 months, plaque-like structures were detectable in representative animals from both groups. Scale bars are indicated in the figure. **(B)** Plaque density in the isocortex and hippocampus, calculated as the number of μXPCT-detected plaque-like structures normalized to the corresponding ROI volume and expressed as plaques/mm³. Each dot represents one animal; horizontal bars and error bars indicate mean ± SEM calculated across all animals within each group. Zero values indicate animals in which no plaque-like structures were detected within the analyzed ROI. Because of the limited number of animals per group, data are presented descriptively and no inferential statistical comparisons were performed.

Plaque burden was quantified within selected isocortical and hippocampal regions of interests (ROIs) by counting morphologically identified plaque structures and normalizing the counts to the corresponding ROI volumes, yielding plaque density (plaques/mm³; **Fig. 4B**). Given the limited number of animals per group (n = 3), plaque-density data were analyzed descriptively, with individual animal values shown together with mean ± SEM and without inferential treatment comparisons.

At 2 months, no μXPCT-detectable plaques were identified in untreated 5xFAD mice within both hippocampus and isocortex, whereas plaques were detected in two of three insulin-treated animals. The available immunofluorescence observations were consistent with these findings, with focal Aβ-positive deposits observed in XPCT plaque-positive samples and no identifiable focal deposits in the examined XPCT plaque-negative samples.

At 6 months, μXPCT-detectable plaques were observed in two of three untreated 5xFAD mice, whereas one animal showed no detectable plaques despite confirmed 5xFAD genotype. Available MOAB-2 sections from this animal likewise showed no identifiable focal Aβ-positive deposits. The basis of this unexpected observation could not be determined from the available data, and the animal was therefore retained in the primary descriptive analysis.

Mean plaque density was higher in insulin-treated than in untreated age-matched 5xFAD mice in both the isocortex and hippocampus. To assess whether this descriptive difference was driven by the plaque-negative untreated 6-month animal, we performed a sensitivity analysis restricted to the two plaque-positive untreated animals. Although exclusion of the plaque-negative animal increased the untreated-group mean, it did not reverse the direction of the treatment-associated difference in either region (**Fig. S5**). Thus, the descriptive pattern was not solely attributable to the zero-value animal. Nevertheless, given the small group size, no definitive conclusion regarding an insulin-dependent increase in amyloid deposition can be drawn. Rather, within the sensitivity of the present μXPCT analysis, insulin treatment was not associated with a detectable reduction in plaque burden. Because the 2-and 6-month cohorts were imaged under different acquisition conditions, direct quantitative comparisons of plaque density between ages should be interpreted cautiously.

### 2.4 Insulin attenuates ileal structural remodeling in 5xFAD mice

To determine whether AD-associated changes and insulin treatment were accompanied by peripheral structural alterations, ileal samples from control and 5xFAD mice, with and without insulin treatment, were examined by μXPCT at 2 and 6 months of age. Reconstructed tomographic volumes enabled label-free three-dimensional visualization of the intestinal mucosa, with villi and crypts clearly resolved within the intact tissue architecture (**Fig. 5A**). Villus length was measured from the villus tip to the villus–crypt junction, and crypt depth from this junction to the crypt base, following their maximal extension in three dimensions. Multiple villi and crypts were measured per animal and summarized at the animal level for statistical analysis.

**Figure 5.**
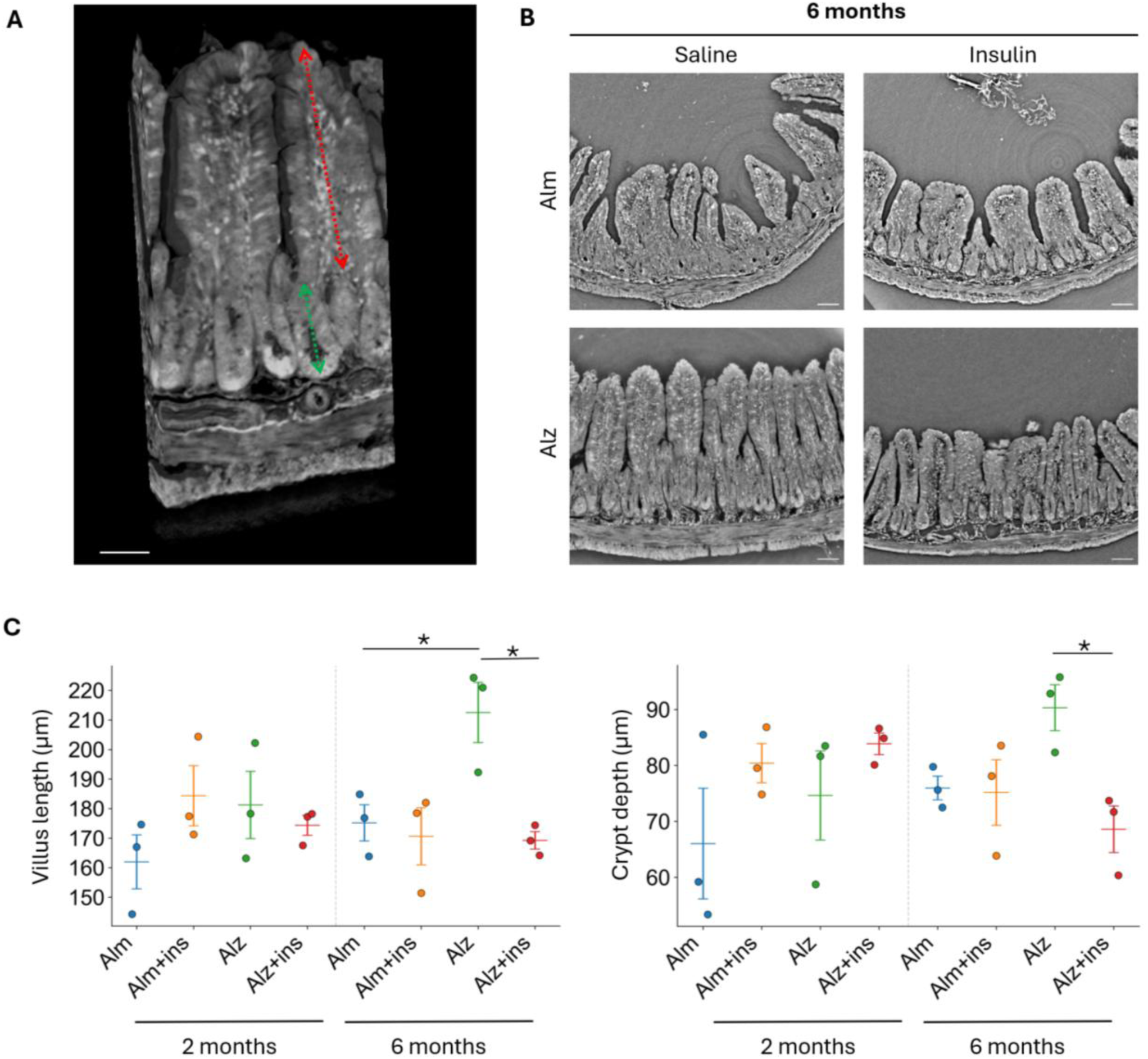
μXPCT-based morphometric analysis of ileal architecture in control and 5xFAD mice. **(A)** Three-dimensional rendering of ileal mucosal architecture reconstructed from a μXPCT volume. The red dashed arrow indicates villus length, measured from the villus tip to the villus–crypt junction, whereas the green dashed arrow indicates crypt depth, measured from the villus–crypt junction to the crypt base. Scale bar: 50 μm. **(B)** Representative μXPCT images of ileum samples from 6-month-old control (Alm) and 5xFAD (Alz) mice, with and without insulin treatment. Scale bars: 50 μm. **(C)** Quantification of villus length and crypt depth across experimental groups at 2 and 6 months of age. Multiple villi and crypts were measured within each sample and summarized at the animal level before group comparison. Each dot represents one animal; horizontal bars and error bars indicate mean ± SEM. Statistical significance was assessed by one-way ANOVA followed by Tukey’s HSD post hoc test; p < 0.05. Asterisks indicate significant pairwise differences.

At 2 months, villus length and crypt depth did not differ significantly among the four experimental groups (**Fig. S6**). In contrast, group differences emerged at 6 months (**Fig. 5B,C**). Untreated 5xFAD mice exhibited significantly greater villus length than untreated control mice (p < 0.05). Villus length was significantly lower in insulin-treated than untreated 5xFAD mice (p < 0.05), with values approaching those observed in controls. Crypt depth showed a related treatment-associated pattern: untreated 5xFAD mice displayed the highest mean crypt depth, although the difference between untreated 5xFAD and control mice did not reach statistical significance. Nevertheless, crypt depth was significantly lower in insulin-treated than untreated 5xFAD mice (p < 0.05). No significant effect of insulin treatment was detected in wild-type mice.

Together, these findings identify an age-dependent ileal structural phenotype in 5xFAD mice, most clearly reflected by villus elongation at 6 months. Insulin treatment was associated with attenuation of villus elongation and lower crypt depth in the 6-month 5xFAD group. Because a significant baseline genotype effect was not established for crypt depth, the insulin-associated decrease in this parameter should not be interpreted as reversal of a disease-associated crypt abnormality or as evidence of functional normalization of the intestine.

### 2.5 Metaproteomics reveals functional remodeling of the gut microbiota

We next asked whether the intestinal structural alterations were accompanied by changes in the fecal microbial proteome. Metaproteomic profiling was therefore performed across control (Alm), insulin-treated control (Alm-ins), 5xFAD (Alz), and insulin-treated 5xFAD (Alz-ins) groups (**Fig. 6**). Peptide-based taxonomic assignments were broadly dominated by *Bacillota* and *Pseudomonadota* across all groups, indicating preservation of the major phylum-level profile (**Fig. 6A**). Differences were more apparent at lower taxonomic levels. *Micrococcales* and *Kitasatosporales* were detected in Alm-ins, Alz, and Alz-ins, while *Herpetosiphonales*- and *Oscillatoriophycideae*-associated assignments were also identified in these groups. These assignments were supported by confidence scores ≥ 0.60 and ≥ 0.76, respectively. *Bacillaceae* and *Anoxybacillaceae* were detected in Alm and Alz-ins with confidence scores ≥ 0.54. These confidence scores reflect support for taxonomic assignment rather than statistical evidence of differential abundance. Functional annotation of the selected peptide set identified hydrolase-associated peptides as the most prominent functional category (**Fig. 6B**). Their relative representation increased from 81% in Alm to 92% in Alm-ins, 96% in Alz, and 93% in Alz-ins (**Table 2**), whereas categories associated with glucose pathways, ATP binding, cation binding, membrane components, and cytosolic components showed more modest differences. Because these functional categories can overlap, these values represent annotations within the selected peptide set rather than mutually exclusive fractions of the total metaproteome. The functional heatmap further revealed distinct abundance patterns among selected peptides (**Fig. 6C**), with several peptides showing lower relative abundance in Alm and higher abundance in Alm-ins, Alz, and Alz-ins. Mapping of the selected peptides to KEGG pathways provided additional functional context for these differences (**Fig. 6D**). Multiple peptides were assigned to proteins involved in ATP turnover and related cellular processes, including F-type ATP synthase subunits ATPA and ATPB, V-type ATPase-associated VATA, as well as DnaK, SecA, RuvB, and MsbA. KEGG mapping included H⁺-transporting two-sector ATPase (EC 7.1.2.2) and inorganic diphosphatase (EC 3.6.1.1). These annotations indicate differential representation of proteins associated with ATP-related processes but do not directly measure ATP production, proton transport, enzymatic activity, or metabolic flux.

**Figure 6.**
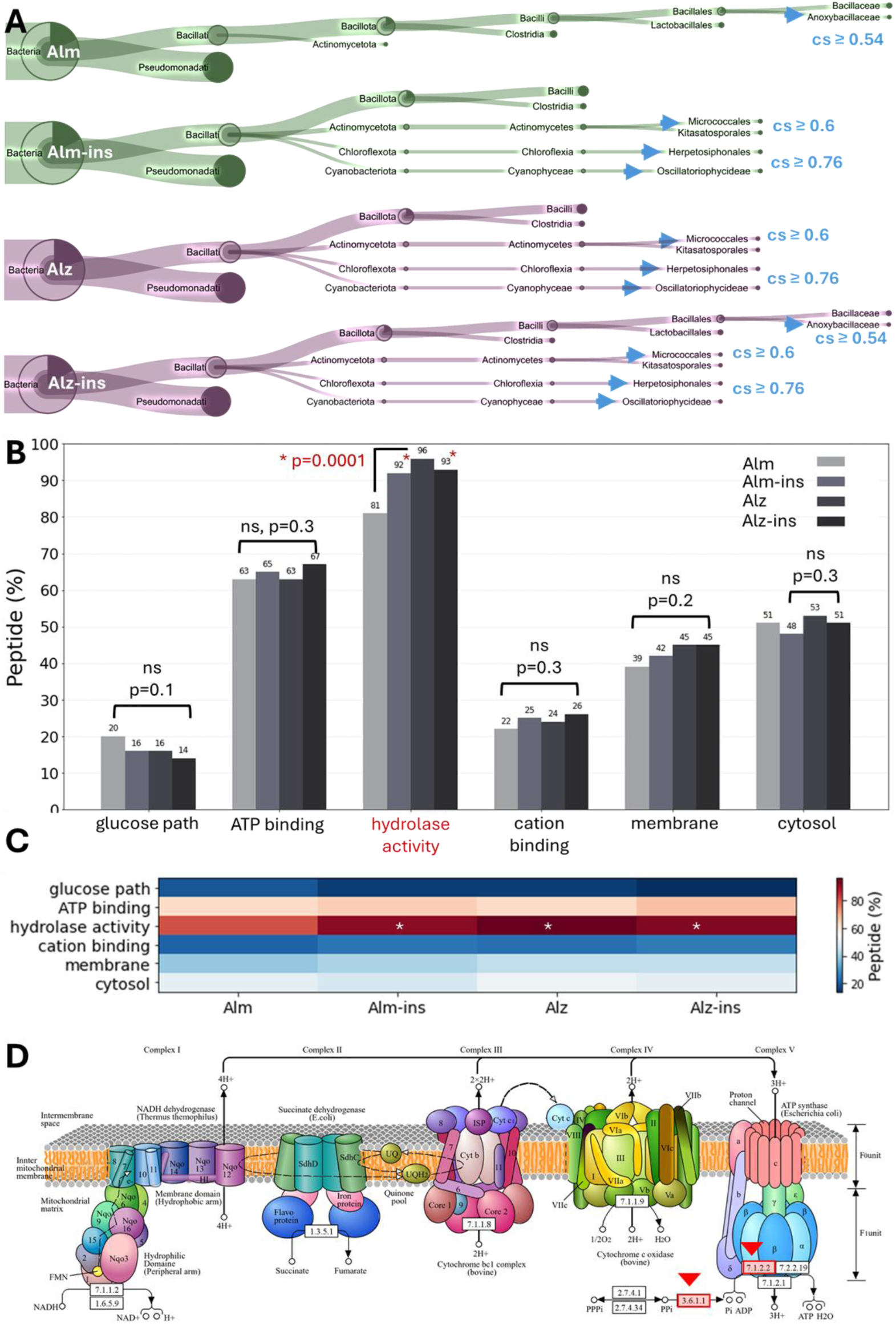
Fecal metaproteomic profiling across experimental groups. (a) Peptide-centric taxonomic profiling showed that the fecal microbiota was predominantly composed of Bacillota and Pseudomonadota in all groups, with additional lower-abundance taxa detected at moderate-to-high confidence (CS). (b) Functional classification of significantly altered peptides identified hydrolase activity as the most prominently enriched category across groups. (c) Heatmap representation of peptide-associated functional categories confirmed the relative enrichment of hydrolase activity, particularly in Alzheimer-associated and insulin-treated conditions. (d) Mapping of altered peptides onto the KEGG oxidative phosphorylation pathway revealed the involvement of ATPase-related proteins linked to microbial energy metabolism.

**Table 2.**
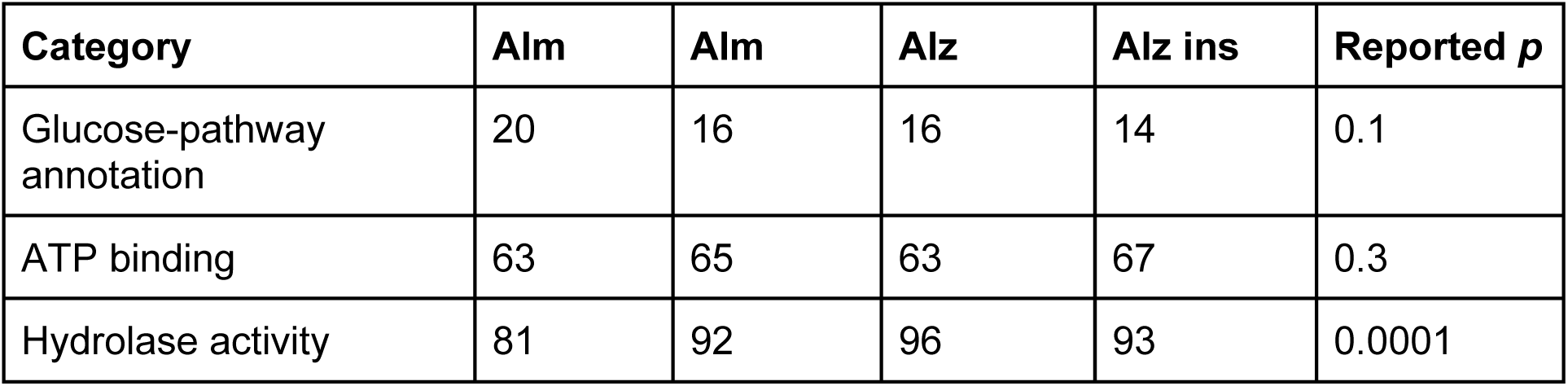
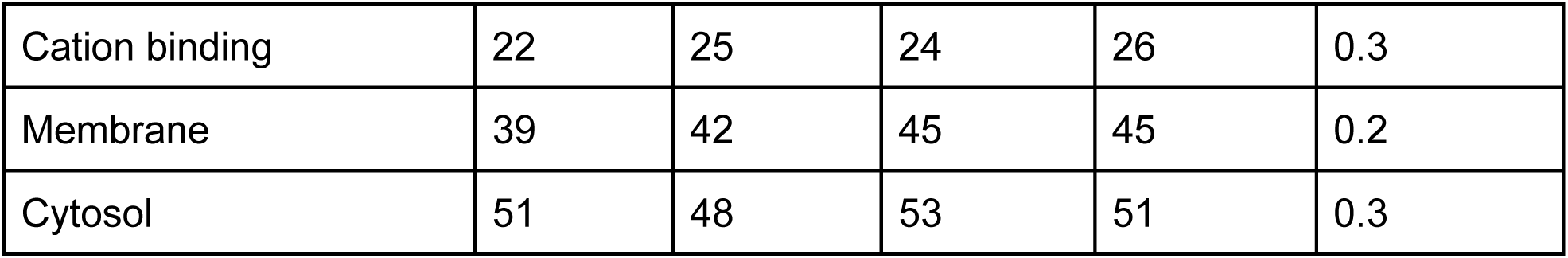
Functional annotation fractions within the selected fecal peptide subset. Values are percentages; overlapping categories need not sum to 100%.

The selected-peptide heatmap contained two broad, opposing patterns (**Fig. S7**). A smaller Alm-high cluster included MscL, some DnaK/ATPB-associated peptides, glucose-6-phosphate isomerase, and elongation factor Tu. A larger cluster relatively low in Alm and high in the other conditions included ATP synthase-associated peptides, elongation factor Tu, triosephosphate isomerase, phosphoglycerate kinase, SecA, DnaK, and ribosomal proteins. Protein-family names in both clusters indicate heterogeneous peptide or variant behavior. Alm-ins, Alz, and Alz-ins shared several abundance trends, but the selected, row-standardized group means do not establish replicate-level clustering or a genotype-independent mechanism. These readouts reveal distinct treatment-associated patterns: detectable brain plaque burden was not reduced, selected ileal structural changes were attenuated in the older 5xFAD cohort, and fecal protein profiles remained different from untreated wild type. Structural attenuation and microbial functional remodeling therefore did not show a uniform shift toward the same reference state.

Collectively, the fecal metaproteomic profiles showed relatively preserved broad taxonomic organization but more pronounced differences in selected functional protein annotations across disease and insulin-treated conditions. Notably, Alm-ins, Alz, and Alz-ins shared several peptide-level patterns relative to untreated controls, suggesting that both disease state and insulin exposure were associated with remodeling of the detectable fecal microbial proteome. These patterns should be interpreted as associations within the measured metaproteomic profiles rather than evidence of a shared underlying metabolic mechanism.

## Discussion

In this study, we integrated XPCT, immunofluorescence and fecal metaproteomics to investigate the effects of intranasal insulin across the brain–gut axis in 5xFAD mice. XPCT provided the main structural readout in brain and ileum, while MOAB-2 immunofluorescence served as a complementary approach to support the identification of Aβ associated pathology. Fecal metaproteomics provided an additional functional layer by characterizing treatment- and disease-associated changes in the microbial proteome.

The main finding was that insulin treatment was associated with distinct responses across biological compartments rather than with a uniform shift toward either a protective or detrimental phenotype. In the brain, insulin-treated 5xFAD mice did not show a reduction in μXPCT-detectable amyloid burden and, descriptively, exhibited higher plaque densities than untreated animals at the analyzed time points (**Fig. 2-4**). In contrast, insulin treatment was associated with reduced villus length and crypt depth relative to untreated 5xFAD mice at 6 months (**Fig. 5**). Metaproteomic profiling further indicated remodeling of the fecal microbial proteome, with functional differences emerging more clearly than broad taxonomic restructuring (**Fig. 6**). Together, these observations indicate that cerebral, intestinal, and microbial responses to intranasal insulin do not necessarily follow the same pattern across the brain–gut axis.

The cerebral findings are particularly relevant in the context of the proposed relationship between insulin signaling and AD. Intranasal insulin has attracted interest because of its potential to influence central insulin signaling while limiting systemic exposure, and previous studies have reported effects on cognition, metabolism, amyloid-related pathways, and neuroinflammatory processes (Ayan and DeMirci, 2023; Freiherr et al., 2013). However, these effects may depend on disease stage, treatment duration, brain region, and the specific pathological endpoint examined; notably, the largest multicenter trial to date found no cognitive or functional benefit in its primary analysis. In the present study, μXPCT detected no plaques in untreated 5xFAD mice at 2 months, whereas plaque-like deposits were detectable in a subset of insulin-treated animals. At 6 months, plaque burden was detectable in both untreated and insulin-treated groups, with higher descriptive mean values in insulin-treated animals in both the isocortex and hippocampus (**Fig. 4**). This pattern persisted when the plaque-negative untreated animal was excluded in the sensitivity analysis. A mechanistic precedent for this direction of effect exists. In neuronal cultures, insulin accelerates the trafficking of APP and Aβ from intracellular compartments toward the plasma membrane, lowering intraneuronal Aβ while increasing its extracellular release (Gasparini et al., 2001). Consistent with this, insulin infusion in cognitively normal older adults acutely raised CSF Aβ42 levels (Watson et al., 2003). Because intraneuronal Aβ precedes extracellular plaque formation in 5xFAD mice (Oakley et al., 2006), a similar insulin-driven shift of Aβ toward the extracellular space could plausibly advance the appearance of detectable deposits, particularly when treatment coincides with the onset of plaque formation. Nevertheless, the small group size and inter-individual variability preclude attributing these differences to a treatment effect. Under the experimental conditions examined, intranasal insulin was therefore not associated with a reduction in μXPCT-detectable amyloid burden.

The multiscale brain XPCT approach provided structural information beyond plaque burden alone. μXPCT enabled label-free visualization of plaque-like deposits within intact paraffin-embedded brains and assessment of their three-dimensional distribution within anatomically defined regions. NanoXPCT further revealed internal heterogeneity and spatial complexity within individual deposits that were only partially resolved at the microscale. MOAB-2 immunofluorescence provided complementary support for the Aβ nature of these structures (**Fig. 3**). Because XPCT and immunofluorescence probe different aspects of the pathology - threedimensional morphology and X-ray contrast on the one hand, and antibody-accessible Aβ associated material in thin sections on the other - the two approaches provide complementary rather than directly interchangeable information.

A different pattern emerged in the intestine (**Fig. 5** and **Fig S6**). At 2 months, villus length and crypt depth were broadly comparable across groups, whereas at 6 months untreated 5xFAD mice showed longer villi and the highest mean crypt depth. Intranasal insulin was associated with lower values for both parameters, with villus length shifting toward control levels and crypt depth showing a similar trend. These findings suggest that insulin attenuated the structural remodeling of the ileal mucosa observed in untreated 5xFAD mice, although the extent of this effect differed between morphometric parameters. The biological significance of these morphological changes remains to be established, but alterations in villus and crypt architecture may reflect changes in epithelial turnover, nutrient handling, mucosal homeostasis, or local inflammatory state (Bonis et al., 2021). Because villus elongation can also represent an adaptive response (Berlin et al., 2019), morphometry alone cannot determine whether the insulin-associated reduction reflects normalization or loss of a compensatory adaptation. Still, this intestinal response occurred in the absence of a corresponding reduction in cerebral amyloid burden, suggesting the possibility that insulin exerts different effects across biological compartments.

The metaproteomic data further support the presence of a compartment-specific response to insulin (**Fig. 6** and **Fig. S7**). Broad phylum-level organization was largely preserved across groups, with *Bacillota* and *Pseudomonadota* remaining dominant, whereas more subtle differences emerged at lower taxonomic levels. In contrast, functional profiling revealed a clearer group-associated signal, with hydrolase activity representing one of the most prominent altered categories and ATPaseand energy metabolism-related proteins contributing substantially to the observed differences. These patterns suggest that disease- and treatment-associated effects may involve changes in microbial metabolic activity and energy-related processes without necessarily producing major restructuring of the dominant microbial phyla. The normalized peptide heatmap also showed that Alm displayed a distinct overall profile, whereas Alm-ins, Alz, and Alz-ins exhibited more similar abundance patterns (**Fig. S7**). This observation raises the possibility that insulin exposure itself contributes to reshaping the fecal microbial proteome, independently of the 5xFAD genotype. A host-side mechanism for such an effect is plausible: mice lacking the insulin receptor specifically in intestinal epithelial cells develop dysbiosis and impaired barrier function, and in diabetic mice insulin treatment alleviated dysbiosis in an intestinal insulin receptor-dependent manner (Gueddouri et al., 2022). Epithelial insulin signaling therefore appears to shape the luminal niche and exogenous insulin reaching the gut could alter microbial activity without requiring the 5XFAD genotype. More broadly, these findings support the view that functional remodeling of the gut microbial community may be more evident than changes in broad taxonomic composition and may therefore provide information that is not captured by taxonomy alone.

An important implication of these findings is that intestinal structural changes and microbial functional remodeling do not necessarily occur in parallel. Although insulin treatment shifted the ileal structural phenotype of 5xFAD mice toward control values, the fecal microbial proteome did not simply revert toward the untreated control state. Instead, insulin-treated control (Alm-ins) and 5xFAD (Alz-ins) groups shared several abundance patterns with the untreated Alzheimer group. This dissociation suggests that changes in tissue morphology should not be assumed to reflect restoration of microbial function. More broadly, the gut–brain axis should not be viewed as a linear pathway in which changes in one compartment necessarily predict changes in another. The relationship is also bidirectional: gut microbiota can themselves regulate brain insulin sensitivity as shown by the transfer of central insulin resistance to germ-free mice by fecal microbiota (Soto et al., 2018). Insulin may instead influence neuronal, epithelial, metabolic, and microbial processes through partially independent mechanisms and timescales. Although the present study does not establish causal relationships among these compartments, it identifies a pattern of differential remodeling that warrants further mechanistic investigation.

Several limitations should be considered. The number of animals per group was limited, particularly for plaque-burden analysis. In addition, the 2- and 6-month cohorts were acquired in separate synchrotron sessions using different acquisition settings, and age-related comparisons were therefore interpreted with appropriate caution. Future studies incorporating additional functional readouts could further clarify the relationships among the observed brain, intestinal, and microbial changes.

Despite these limitations, the integration of XPCT, immunofluorescence, and fecal metaproteomics provided complementary structural and molecular information across the brain–gut axis. The data indicates that intranasal insulin was associated with persistent cerebral amyloid pathology, without evidence of amyloid-related benefit, reduced villus length and crypt depth in 5xFAD ileum, and continued functional reorganization of the fecal microbial proteome. Rather than supporting a uniform effect of insulin across tissues, these findings point to distinct responses across cerebral, intestinal, and microbial compartments. Larger studies incorporating direct functional and mechanistic measurements will be required to determine how these responses are related and whether they influence disease progression or treatment response.

## Conclusions

Intranasal insulin was associated with distinct structural and molecular patterns across the brain–gut axis in this exploratory 5xFAD study. XPCT-detectable plaque burden was not reduced, ileal villus and crypt dimensions were lower in the treated older cohort, and fecal peptide profiles remained different from untreated wild type. These findings distinguish structural attenuation from microbial functional reversion and plaque visibility from total amyloid quantity. Multiscale XPCT, independent molecular labeling, and metaproteomics provide complementary tools for evaluating such differences. Larger, fully documented studies are needed to establish reproducibility, mechanism, and relevance to disease progression or treatment response.

## Methods

### 3.1. Samples

The study used 5XFAD transgenic mice and their littermates obtained from Jackson Laboratory (Bar Harbor, Maine, USA). Analyses were performed on excised brain and intestinal tissues from 24 mice in total, divided into four groups. The control group (Alm) included 6 mice, with 3 animals analyzed at 2 months and 3 at 6 months. The insulin-treated control group (Alm-ins) also included 6 mice, with 3 analyzed at 2 months and 3 at 6 months. The Alzheimer’s group (Alz) consisted of 6 mice, again with 3 analyzed at 2 months and 3 at 6 months. Finally, the insulin-treated Alzheimer’s group (Alz-ins) included 6 mice, with the same age distribution. Table X summarizes the animal models and the number of samples analyzed in this study.

The 5xFAD mouse model (#034848-JAX) is widely used in Alzheimer’s research because it develops robust amyloid-β pathology at an early age, with plaque formation beginning around 2-4 months and progressing over time (Oakley et al., 2006). These mice carry five familial Alzheimer’s mutations in *APP* and *PSEN1*, which drive rapid accumulation of amyloid-β, neuroinflammation, and neuronal dysfunction. The model also shows a strong and reproducible plaque burden across key brain regions, including the cortex and hippocampus, making it especially useful for investigating amyloid-related mechanisms. For this reason, 5xFAD mice are well-suited for insulin studies, as they allow direct assessment of how insulin treatment may affect plaque development, disease progression, and related pathological changes in such a well-characterized amyloid-dominant system.

The mice were supplied by the Istanbul Medipol University Health Sciences and Technologies Research Institute (SABITA). Environmental enrichment was intentionally omitted because it has been reported to worsen Alzheimer’s-like pathology in murine models. Outside the specified treatment intervals, the animals were not exposed to additional pharmacological agents and did not undergo behavioral testing. They were housed in standard mouse cages with sawdust bedding and had free access to food and water. The SABITA animal facility maintained conventional laboratory conditions, with a temperature of approximately 20 ± 2°C, relative humidity of 60%, and a 12-hour light--dark cycle from 7:00 a.m. to 7:00 p.m.

#### 3.1.1 Genotyping, polymerase chain reaction, agarose gel electrophoresis, and treatment

Genotyping was performed using tissue samples collected from 28–30-day-old mice prior to weaning while still under maternal care. Following light anesthesia with isoflurane, identification ear tags were applied to the right ear, and approximately 1 mm² of tissue was obtained from the same ear using a punch tool. The collected tissue was transferred into 0.2 ml PCR tubes containing 75 μl of alkaline lysis buffer (25 mM NaOH/EDTA) and incubated at 98°C for 1 hour using a T100™ Thermal Cycler. Subsequently, 75 μl of neutralization buffer (40 mM Tris/HCl) was added. After pipetting and vortexing, the samples were centrifuged at 4000 rpm for 4 minutes. The resulting supernatant was then carefully transferred into 0.5 ml tubes and stored at −20°C until further use.

PCR reactions were prepared by combining 0.5 μM of both forward and transgenic primers, 200 μM dNTP mixture, 1.25 units of Taq DNA polymerase, template DNA, and ddH₂O in 1× reaction buffer to a final volume of 12 μl. A negative control lacking template DNA was included in each run. The primer sequences used were as follows: forward primer (5’-GAA AAG ATA AGG CCT TCA AGT GCT-3’), internal positive control reverse primer (5’-GAT CTT TCT GCA GTA TCT TCC TTC-3’), and transgenic primer (5’-ATC GAC CGG TAA TGC AGG CAA-3’). The prepared PCR mixtures were then amplified using the specified protocol in a T100™ Thermal Cycler.

Agarose gel electrophoresis was performed by dissolving 2 g of agarose in 100 ml of 1× TAE buffer (40 mM Tris, 20 mM glacial acetic acid, 1 mM EDTA, pH 7.6) using a microwave until the solution became clear. After cooling slightly, 5 μl of SafeView was added, and the solution was poured into an electrophoresis cassette with loading combs. Once solidified, the gel was placed into an electrophoresis tank containing 1× TAE buffer. Each well was loaded with 9 μl of PCR product mixed with 1 μl of loading dye, and 5 μl of DNA marker was loaded into the first well. Electrophoresis was performed at 100 V for 20 minutes. Following separation, bands were visualized using a Bio-Rad ChemiDoc imaging system, and the resulting band patterns were analyzed to determine the transgenic status of the mice.

Mice in the control and Alzheimer’s subgroups (Alm, Alz) received intranasal physiological saline daily, whereas mice in the insulin-treated subgroups (Alm-ins and Alz-ins) received intranasal insulin on the same protocol. In the insulin-treated groups, U-100 insulin was administered for 5 weeks at a dose of 1 U per mouse. Each administration was delivered in a total volume of 10 μL, corresponding to approximately 5 μL per nostril.

### 3.3 Tissue preparation for X-ray imaging

Mice were euthanized by CO₂ inhalation. Brains and ileum samples were dissected and fixed for 24 h at 4 °C in 4% paraformaldehyde (PFA) in phosphate-buffered saline (PBS). Following fixation, samples were dehydrated through a graded ethanol series (70%, 95%, and 100%), cleared with xylene and subsequently embedded in paraffin wax. Paraffin embedding was performed while preserving the overall tissue morphology, allowing the specimens to be imaged intact by micro-X-ray phase-contrast tomography and subsequently punched to obtain 500-µm-diameter cylindrical biopsies for nanoXPCT analysis. No exogenous contrast agent or heavy-metal staining was used at any stage prior to X-ray imaging; therefore, image contrast relied on the intrinsic differences in X-ray attenuation and refractive properties of the tissues.

### 3.4. Micro X-ray phase contrast tomography (μXPCT)

μXPCT measurements were performed in two independent experimental sessions. The first experiment was carried out at the SYRMEP beamline of Elettra Synchrotron (Trieste, Italy) on samples from 2-month-old mice, whereas the second experiment was performed at the ID19 beamline of the European Synchrotron Radiation Facility (ESRF, Grenoble, France) on samples from 6-month-old mice. For all μXPCT measurements, intact paraffin-embedded specimens were scanned to reconstruct three-dimensional volumes of the entire organs or selected regions of interest (ROIs). The embedded brain samples had approximate dimensions of 0.8 × 0.5 × 1.0 cm³ (width × depth × height), while ileum specimens were smaller, with typical dimensions of approximately 0.2 × 0.2 × 0.8 cm³.

At the SYRMEP beamline, μXPCT measurements were performed using a pink beam with an energy distribution centered at approximately 16.7 keV. Data were collected at two spatial resolutions. Whole-brain samples were scanned with an effective pixel size of 2.2 μm, using a sample-to-detector distance of 200 mm and an exposure time of 50 ms per projection. Higher-resolution scans were acquired on selected brain ROIs and at different positions along the ileum samples, using an effective pixel size of 1.2 μm, a sample-to-detector distance of 120 mm, and an exposure time of 50 ms per projection angle. Tomographic datasets consisted of 3600 projections.

At the ID19 beamline, μXPCT measurements were performed using a pink beam with an energy of approximately 22 keV. Samples were scanned at two spatial resolutions. Scans with an effective pixel size of 0.6 μm were acquired using a sample-to-detector distance of 200 mm and an exposure time of 0.1 s per projection. Higher-resolution scans with an effective pixel size of 0.3 μm were acquired using a sample-to-detector distance of 80 mm and an exposure time of 0.05 s per projection. For each scan, 4500 projections were collected.

For both experimental sessions, tomographic datasets were acquired in extended field-of-view (FOV) mode, a scanning approach that increases the effective horizontal coverage of the detector.

In this configuration, the rotation axis is displaced toward one side of the detector FOV, and projections spanning 360° are recorded with a width corresponding to the detector size. The sinograms were subsequently stitched prior to tomographic reconstruction. Preprocessing of the projections, phase retrieval, and tomographic reconstruction were performed using beamline-specific reconstruction pipelines. Phase retrieval was performed using Paganin’s algorithm (Paganin et al., 2002). A single projection image per angle was collected in free-space propagation mode at a defocused position corresponding approximately to the Fresnel distance, where phase contrast is maximized. Under these conditions, the radiation wavelength and the sample composition satisfied the algorithm’s assumptions. This approach enables simultaneous retrieval of the phase and absorption components of the transmitted wave through their ratio. Three-dimensional volumes were reconstructed from the phase-retrieved projections using the Filtered Back Projection algorithm.

### 3.5. Nano X-ray phase contrast tomography (nXPCT)

Guided by microXPCT images, 500-μm biopsies were punched out from the identified ROI of brains and imaged at the Nano-Imaging ID16A beamline of the ESRF (Grenoble, France) with an effective pixel size of 80 nm. A pair of multilayer-coated Kirkpatrick-Baez optics was used to focus the X-rays (∼30 nm) at 33.35 keV. The sample was put in the divergent beam downstream of the focus to produce magnified phase-contrast images. The projection geometry also allows zooming into specific regions of a large sample by combining scans with different magnifications and FOV (Bartels et al., 2015; Mokso et al., 2007). By measuring the Fresnel diffraction patterns at four different effective propagation distances, the phase maps of the sample can be retrieved via holographic reconstruction, with the so-called phase-retrieval procedure (Cloetens et al., 1999) implemented in GNU Octave. Magnified radiographs were recorded onto an X-ray detector using a scientific CMOS camera from XIMEA GmbH, with a GSENSE6060 sensor from Gpixel, 6144 × 6144 pixels with each 10 μm edge length. For a single tomography scan, 2000 projections were acquired with an exposure time of 0.2 s. Tomography scans at four different focus-to-sample distances (i.e., four different Fresnel numbers) were collected to complete a single holotomography scan. The tomographic reconstruction was obtained with the ESRF PyHST software package (Paleo et al., 2024).

### 3.6. XPCT image analysis and quantification

Image processing and segmentation were performed using ImageJ. Prior to analysis, all datasets were preprocessed to reduce artifacts arising from data collection parameters and tomographic reconstruction. In *μ*XPCT datasets, grayscale intensity is proportional to electron density, with brighter pixels corresponding to higher-density structures and darker pixels to lower-density regions. For nanoXPCT images, where specified, the grayscale was inverted for visualization purposes to improve the visual perception of internal structural features; this display adjustment did not affect the underlying image data or any quantitative analysis.

#### 3.6.1 Amyloid-β plaque quantification

Amyloid-β plaques were manually identified and counted in μXPCT brain images within predefined regions of interest (ROIs). ROIs were assigned to two anatomical areas, namely the hippocampus and the isocortex, and plaque quantification was performed separately for each region. Plaque distribution was expressed as plaque density, calculated as the number of detected plaques divided by the corresponding ROI volume, and reported as plaques/mm³. The reliability of plaque recognition was assessed by independent counting performed by two different operators. Because of the limited number of animals per group, plaque-density data were analyzed descriptively and no inferential statistical comparisons were performed between experimental groups. All animals with confirmed group assignment were included in the primary descriptive analysis, including animals in which no plaques were detected by μXPCT. At 6 months, one untreated 5xFAD mouse showed no μXPCT-detectable plaques despite confirmed genotype. To evaluate the influence of this plaque-negative animal on the group-level trend, a sensitivity analysis was additionally performed for the untreated 5xFAD group by recalculating the descriptive mean using only the plaque-positive animals. Results from both the complete group and the plaque-positive subset were reported to provide a transparent assessment of the robustness of the observed trend.

#### 3.6.2 Intestinal morphometric analysis

Morphometric analysis of the ileum was performed on XPCT datasets to quantify morphometric parameters of the intestinal mucosa. Villus length and crypt depth were measured from reconstructed 3D volumes, enabling accurate identification of the maximum z-axis extension. To minimize measurement bias, villus length was measured at the point of maximum elongation, and crypt depth was determined using the same criterion. Approximately 200 villi and 150 crypts per sample were analyzed. Measurements were performed independently by multiple observers in a blinded manner to improve reproducibility.

### 3.7. Immunofluorescence

Paraffin-embedded brain sections (7 µm) were baked at 60°C for 1 hour. The sections were then deparaffinized in xylene (twice, 5 minutes each) and rehydrated through a graded ethanol series (100%, 100%, 95%, and 70%; 3 minutes each), followed by rinsing in distilled water. Heat-induced epitope retrieval was performed in 1X citrate buffer (pH 6.0). The slides were heated in a microwave at 800 W for 2.5 minutes, followed by 180 W for 15 minutes, and then allowed to cool at room temperature for 30 minutes. After washing with phosphate-buffered saline (PBS), the sections were blocked and permeabilized for 1 hour at room temperature in a blocking buffer containing 3% (w/v) bovine serum albumin (BSA), 0.3% (w/v) sodium azide, 1% (v/v) normal goat serum, 0.1 M glycine, and 0.1% (v/v) Triton X-100 in PBS. Then, the sections were incubated overnight at 4°C in a humidified chamber with mouse monoclonal anti-amyloid-β antibody MOAB-2 (Abcam, ab126649; 1:600). The primary antibody was prepared in an antibody dilution buffer containing 3% (w/v) BSA, 0.3% (w/v) sodium azide, 1% (v/v) normal goat serum, 0.1 M glycine, and 0.1% (v/v) Tween-20 in PBS. The following day, the sections were washed three times with PBS for 5 minutes each and incubated with Alexa Fluor Plus 488 secondary antibody (Invitrogen, A32723; 1:1500), prepared in the same antibody dilution buffer, for 1 hour at room temperature in the dark. Cell nuclei were then counterstained with DAPI (1:1000) for 5 minutes. After two final PBS washes of 3 minutes each, the slides were coverslipped using Fluoromount™ aqueous mounting medium (Sigma-Aldrich, F4680) and stored at 4°C in the dark until imaging. Primary-antibody-omission controls were processed in parallel by replacing MOAB-2 with the antibody dilution buffer while keeping all subsequent steps identical. These controls were used to assess primary-antibody-independent background and nonspecific binding of the secondary antibody. Fluorescence images were acquired using a Zeiss Axio Zoom.V16 microscope equipped with a PlanNeoFluar Z 1.0x/0.25 objective and an AxioCam HR R3 camera at 11.2x zoom using ZEN Blue software. Alexa Fluor 488 and DAPI were imaged using the corresponding filter sets. Although no red fluorophore was used, an additional red reference channel (excitation, 559–585 nm; emission, 600–690 nm) was acquired to qualitatively assess broad-spectrum tissue autofluorescence. Channel-specific acquisition settings were kept unchanged across MOAB-2-stained and primary-antibody-omission control sections.

### 3.8. Metaproteomics analysis, data processing, and refinement

Metaproteomics analysis has been performed as mentioned in our previous work (Ayan et al., 2023). Briefly, we collected intestinal samples from mice and stored them at −80 °C until analysis. After thawing, we isolated bacterial material from fecal samples by vortexing in phosphate-buffered saline (PBS), shaking for 45 minutes, and centrifuging three times. The supernatants were combined and centrifuged again at 20,000×g. We extracted proteins from 25 mg of the resulting pellet using a buffer of 2% SDS, 100 mM DTT, and 20 mM Tris-HCl (pH 8.8), followed by heating, freezing, reheating, sonication, and centrifugation. The supernatants were stored at −80 °C. For digestion, we followed the FASP protocol (Wiśniewski, 2019), loading 100 µg of protein onto 30-kDa spin columns in 8 M urea/Tris-HCl. After washing and reducing with DTT, the samples were alkylated in the dark with iodoacetamide and digested with trypsin for 18 hours at 37 °C. We eluted the peptides, concentrated them under vacuum, quantified using a colorimetric peptide assay, and injected 5 µL for LC–MS/MS runs. Using an ACQUITY UPLC M-Class system and a SYNAPT Xevo G2-XS mass spectrometer, we performed Nano LC–MS/MS analysis. Peptides were separated on a C18 column at 55 °C, and data were acquired in SONAR mode. The raw data were processed with Progenesis QI for Proteomics against a UniProt database. Proteins were considered differentially expressed with ANOVA p<0.05, q<0.05, more than 2 unique peptides, and a fold change of ≥ 1.3.

Data analyses were further refined using the Unipept suite (Verschaffelt et al., 2021), which enables peptide-centric taxonomic and functional profiling of microbial communities. Identified tryptic peptides were matched against the UniProtKB database using exact sequence matching. Taxonomic assignments were obtained using the Lowest Common Ancestor (LCA) approach, which integrates all taxonomic annotations associated with matched protein entries. The resulting microbial composition was visualized using the Unipept treeview interface. Functional annotation was conducted by mapping peptides to Gene Ontology (GO) terms derived from UniProt entries, allowing classification into biological process, molecular function, and cellular component categories. Statistical evaluation of taxonomic and functional differences between groups was performed using the Unipept peptonizer tool (by confidence score), which aggregates peptide-level information to assess differential abundance and significance across conditions.

To visualize differential peptide abundance, statistically significant peptides (p < 0.05) identified by one-way ANOVA were selected for heatmap analysis. Quantitative abundance values (group means) were extracted using custom Python scripts, then log2-transformed (log2(x + 1)) to address dynamic range and zero inflation in metaproteomic data. Row-wise z-score normalization (z=(x−μ)/σ) was applied (Easterly et al., 2019), excluding peptides with zero variance. Missing values were removed before transformation. Hierarchical clustering used Euclidean distance and average linkage for peptides, with color intensities indicating z-scores, where red indicates enrichment and blue indicates depletion of peptide abundance across conditions.

### 3.9. Statistics

For statistical analyses, animals were considered the independent experimental units. For each experimental group, n represents the number of animals included in the analysis. When multiple technical or object-level measurements were obtained from the same animal, including plaques, villi, crypts, or other XPCT-derived measurements, values were summarized at the animal level prior to group comparisons. Amyloid-β plaque-density data were analyzed descriptively because of the limited number of animals per group; therefore, no inferential statistical comparisons were performed for plaque burden. For the 6-month untreated 5xFAD group, an additional sensitivity analysis was performed by recalculating the descriptive group mean using only plaque-positive animals. Metaproteomic statistical analyses were performed using MedCalc software (MedCalc Software Ltd., Belgium). Following functional annotation using the *Unipept* platform, differences between experimental groups were assessed relative to the control group (*Alm*). For pairwise comparisons of proportions, a one-sample or two-sample test for proportions (z-test) was applied as appropriate. For comparisons involving multiple groups, one-way analysis of variance (ANOVA) was used, followed by post hoc pairwise comparisons. *p*-values < 0.05 were considered statistically significant. Data are presented as mean values ± standard deviation unless otherwise stated.

## Author contributions

E.A., F.P., M.I.A., and A.C. designed the research. E.A., F.P., Z.D., A.A.K., I.B., L.M., E.L., G.B., S.B., and K.K. performed experiments. E.A., F.P., G.C. and I.B. analyzed and refined the data. A.K., E.E., G.O., M.I.A., and A.C. provided resources and supervised the research. E.A. and F.P. wrote the manuscript.

## Acknowledgement

We acknowledge Elettra Sincrotrone Trieste for providing access to synchrotron radiation facilities number 20245455, and we thank Elettra staff for assistance in using beamline SYRMEP. We acknowledge the European Synchrotron Radiation Facility (<u>ESRF</u>) for the provision of synchrotron radiation facilities under proposal ID MD1434on beamline ID16A and LS3461 on beamline ID19. We thank Murielle Salomè (ID16A) and Marta Majkut (ID19) for assistance and support during the beamtime. We also thank Ahmet Tarık Baykal for support during metaproteomic analysis.

---

**Figure S1.**
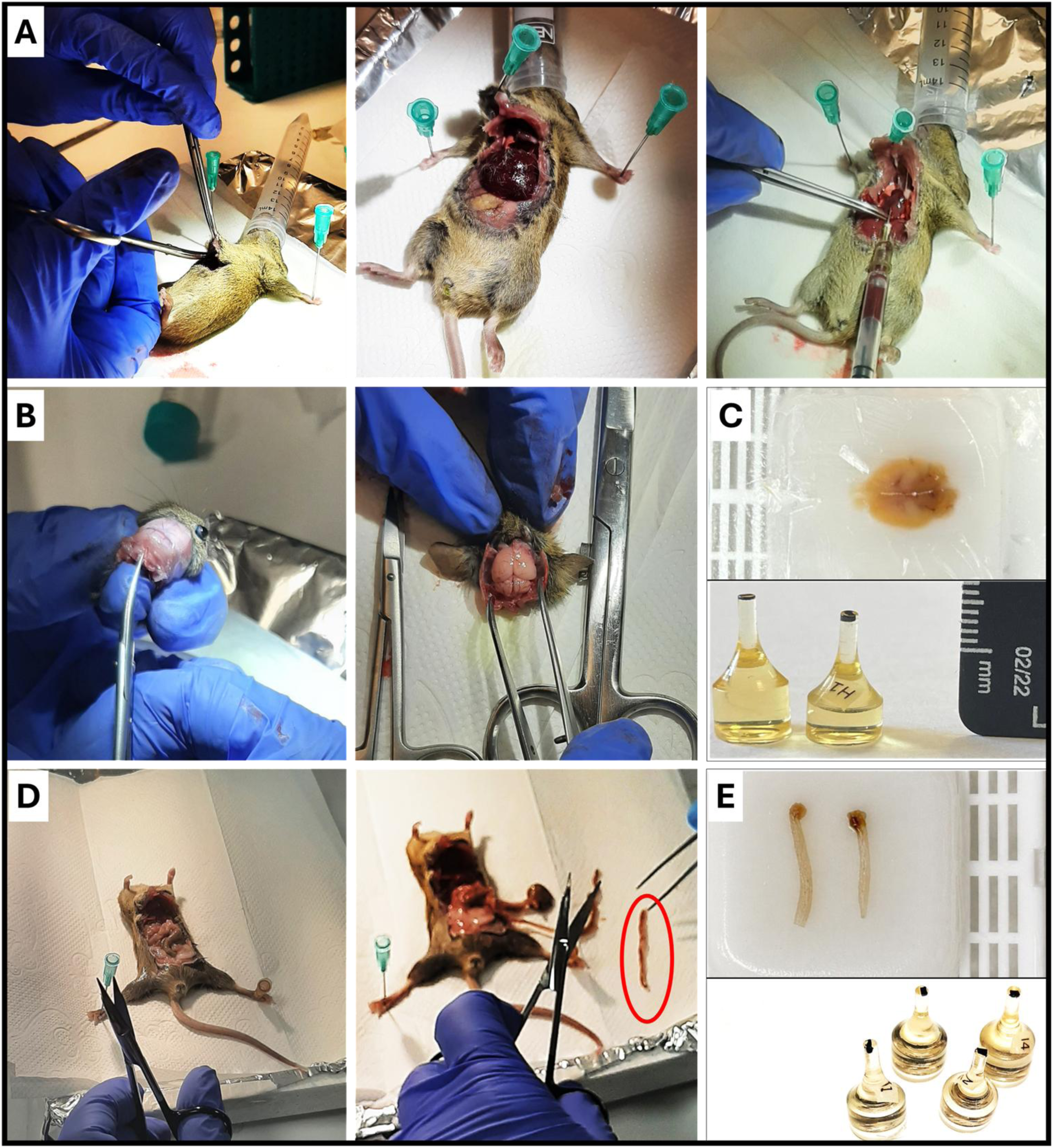
Experimental workflow for dissection and XPCT sample preparation. (A) Representative image of terminal blood sampling by cardiac puncture. (B) Brain dissection during tissue collection. (C) Brain sample prepared for imaging, including the paraffin-embedded whole brain (top) and epoxy-embedded tissue specimen (bottom). (D) Ileum dissection from the abdominal cavity; the excised intestinal segment is highlighted in red coloring. (E) Ileum specimens prepared for imaging, including paraffin-embedded samples (top) and epoxy-embedded tissue specimens (bottom).

**Figure S2.**
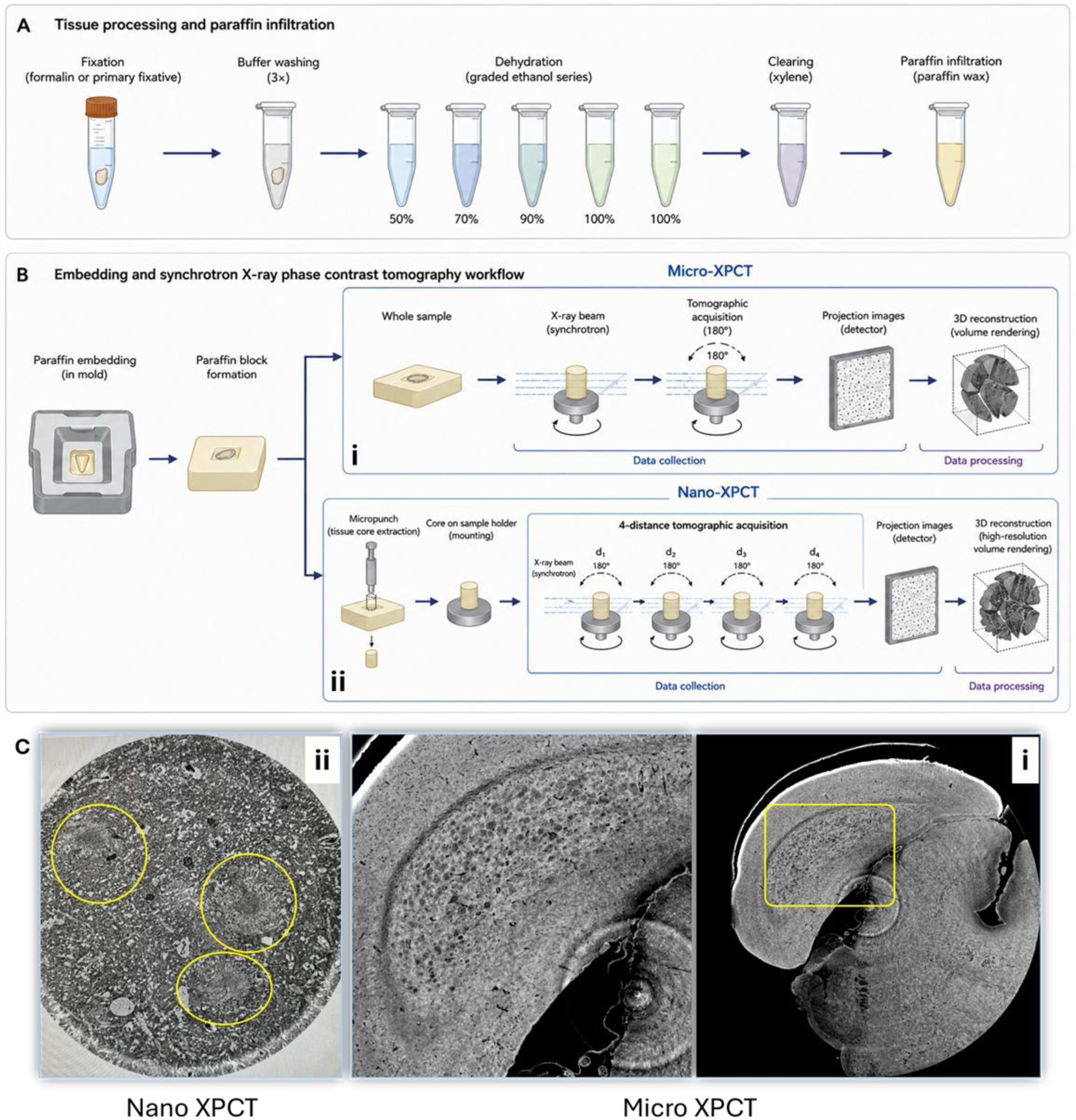
Multiscale XPCT workflow for micro- and nano-scale imaging of paraffin-embedded tissues. (A) Schematic overview of sample preparation for XPCT imaging. Following fixation, samples were dehydrated through a graded ethanol series, treated with propylene oxide, and embedded in paraffin wax. (B) Multiscale XPCT workflow. For μXPCT, intact paraffin-embedded specimens were imaged using extended-field-of-view acquisition over 360°, followed by phase retrieval and tomographic reconstruction. For nanoXPCT, μXPCT datasets were used to guide the extraction of 500-µm-diameter cylindrical biopsies from selected brain regions. Samples were subsequently imaged using four-distance holotomographic acquisition and reconstructed at high spatial resolution. (C) Representative μXPCT and nanoXPCT reconstructions illustrating the transition from whole-specimen imaging to high-resolution characterization of selected regions.

**Figure S3.**
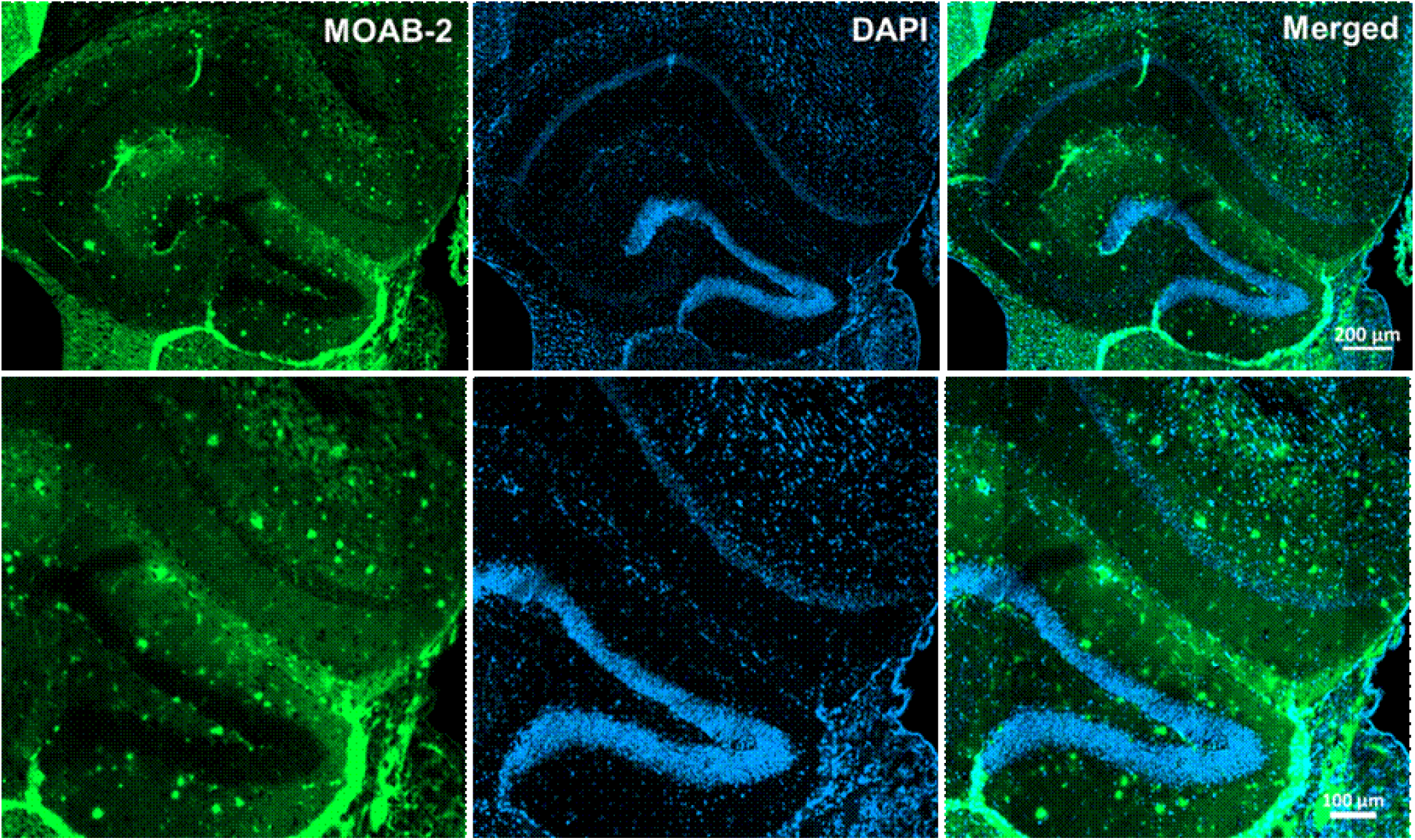
MOAB-2 immunofluorescence of amyloid-β pathology in a 6-month-old insulin-treated 5xFAD mouse. Representative immunofluorescence images of the hippocampal region from a 6-month-old insulin-treated 5xFAD mouse (Alz-ins). The top row shows a panoramic view of the hippocampal formation, while the bottom row shows a higher-magnification view of the selected region. Aβ-associated deposits are labeled with MOAB-2 (green), and cell nuclei are counterstained with DAPI (blue); merged images are shown in the right panels. Scale bars: 200 μm (top row) and 100 μm (bottom row).

**Figure S4.**
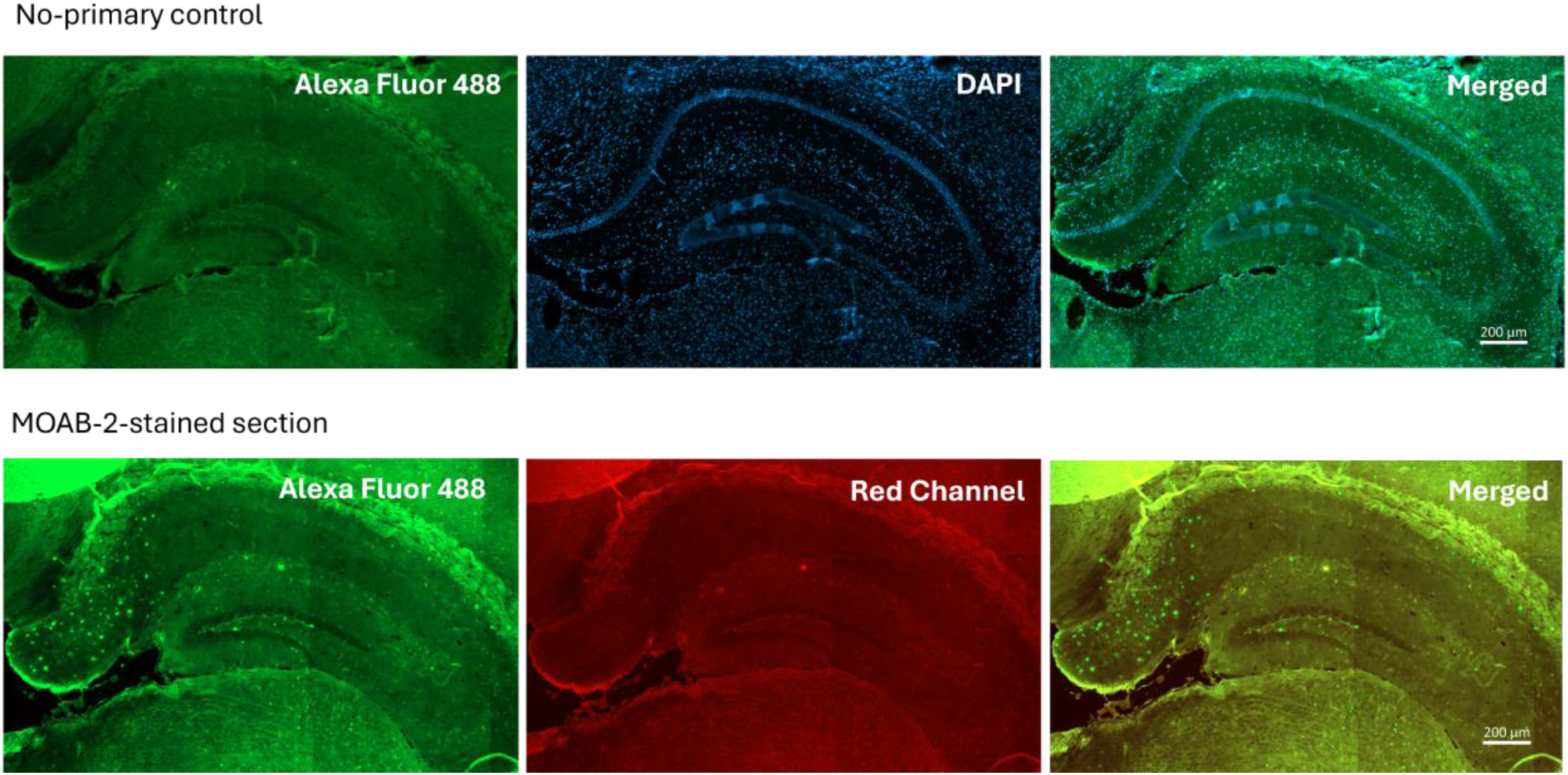
Assessment of background fluorescence and tissue autofluorescence in MOAB-2 immunostaining. The top row shows hippocampal sections processed without the primary antibody and incubated with the Alexa Fluor Plus 488 secondary antibody. Alexa Fluor 488 signals detected under these conditions were considered primary-antibody-independent background. The bottom row shows a MOAB-2-stained hippocampal section imaged in the Alexa Fluor 488 and red reference channels. Structures visible in both channels were considered potentially autofluorescent, whereas additional discrete Alexa Fluor 488-positive puncta without a corresponding red-channel signal were considered consistent with Aβ-associated immunoreactivity. Scale bars: 200 µm.

**Figure S5.**
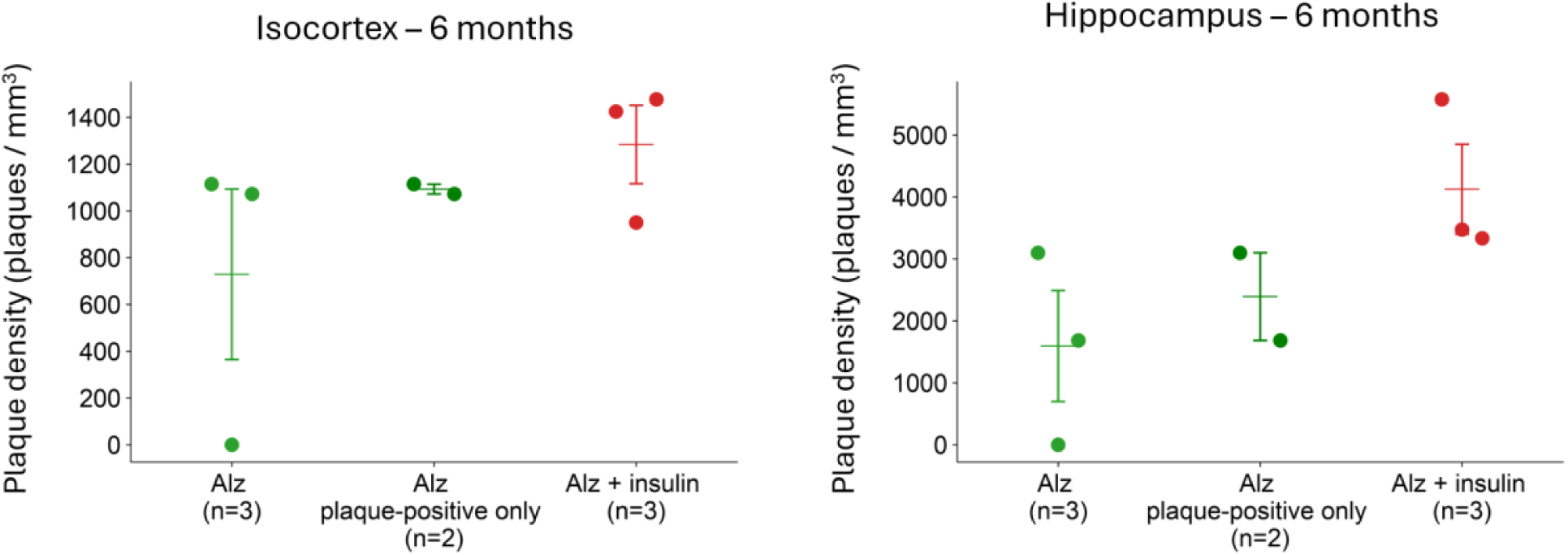
Sensitivity analysis of plaque density in 6-month-old 5xFAD mice. Plaque density in the isocortex and hippocampus is shown for untreated 5xFAD mice including all animals (Alz, n = 3), the same group restricted to animals with μXPCT-detectable plaques (Alz, plaque-positive only, n = 2), and insulin-treated 5xFAD mice (Alz + insulin, n = 3). Individual animal values are shown together with mean ± SEM. The analysis was performed to assess the influence of the plaque-negative untreated animal on the descriptive group mean. Mean plaque density remained lower in the plaque-positive untreated subset than in insulin-treated 5xFAD mice in both analyzed regions.

**Figure S6.**
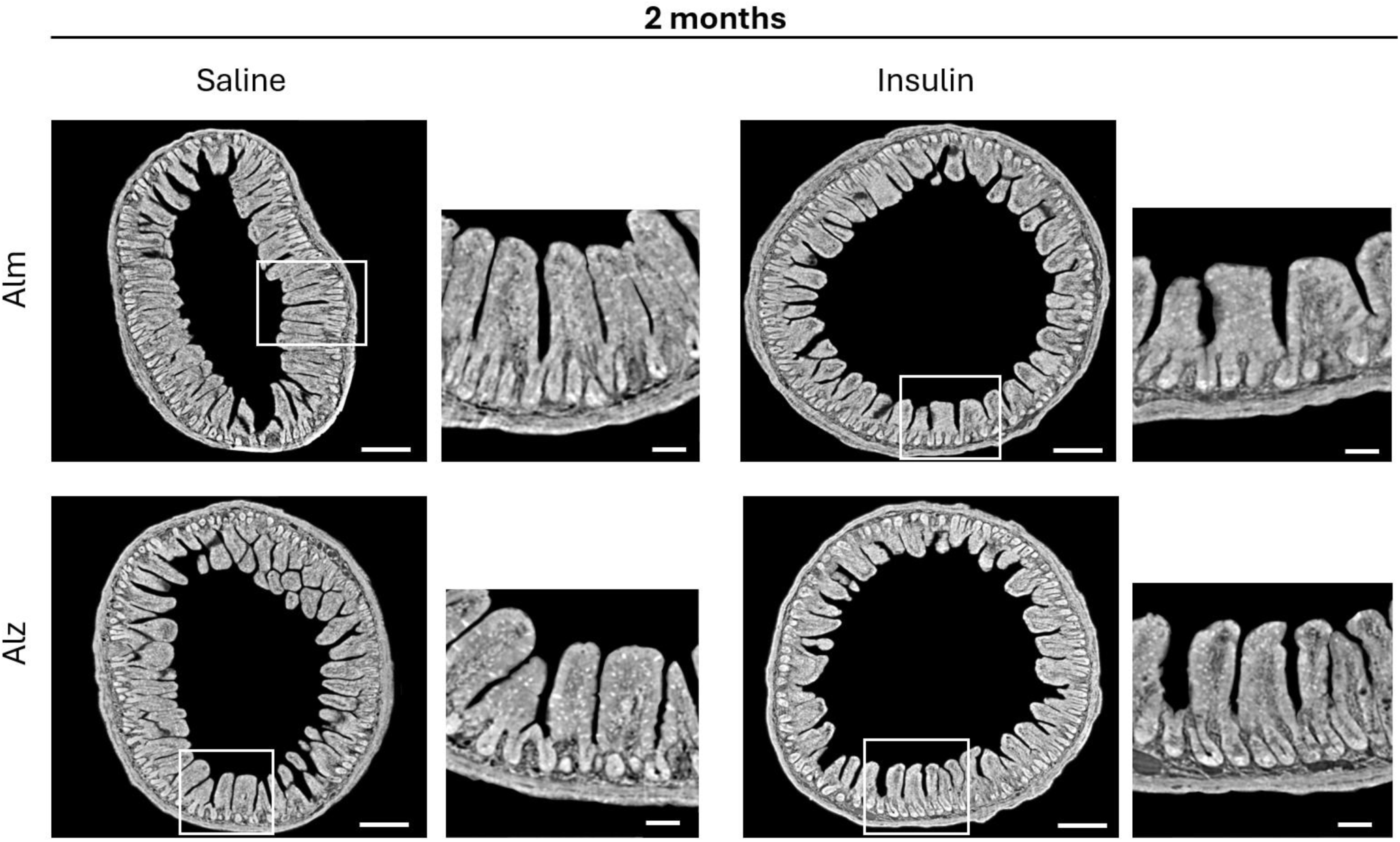
μXPCT visualization of ileal architecture in 2-month-old mice. Representative μXPCT images of ileum samples from control mice (Alm) and 5xFAD mice (Alz), with and without insulin treatment, at 2 months of age. Whole cross-sections are shown together with magnified views of selected mucosal regions, highlighting villus and crypt architecture. White boxes indicate the regions selected for magnification. Scale bars: 200 μm for whole cross-sections and 50 μm for magnified views.

**Figure S7.**
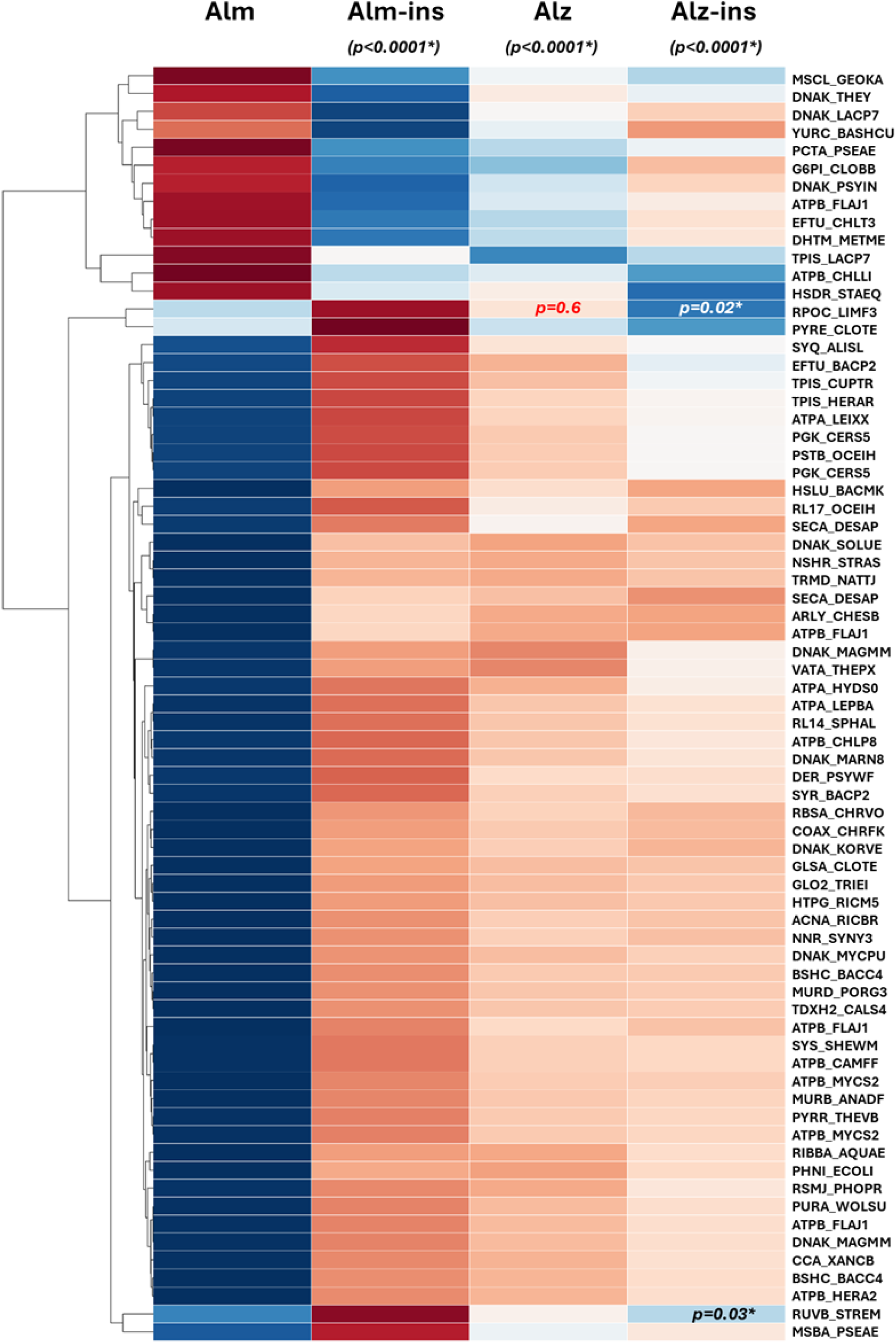
Row-wise normalized peptide abundances are shown for Alm, Alm-ins, Alz, and Alz-ins. The Alm exhibited a distinct peptide profile, whereas Alm-ins, Alz, and Alz-ins showed more similar abundance patterns. Two major peptide clusters with opposite trends were evident across groups. Most statistically significant peptides showed strong significance (p < 0.0001), and many were associated with ATPase- and energy-metabolism functions, consistent with functional remodeling of the gut microbial proteome.

**Table 3.** Experimental animal projection.

| Model | Age | Nu (n) | Description |
| --- | --- | --- | --- |
| C57BL/6 | 8 weeks (2 months) | 3 | 1 U per day of U-100 insulin (~5 µL per nostril) has been administered over 5 weeks. |
| C57BL/6 (INS) | 8 weeks (2 months) | 3 |  |
| C57BL/6 | 24 weeks (6 months) | 3 |  |
| C57BL/6 (INS) | 24 weeks (6 months) | 3 |  |
| 5XFAD | 8 weeks (2 months) | 3 |  |
| 5XFAD (INS) | 8 weeks (2 months) | 3 |  |
| 5XFAD | 24 weeks (6 months) | 3 |  |
| 5XFAD (INS) | 24 weeks (6 months) | 3 |  |

